# Diet-Related Molecular Evolution Differs among Vertivores, Invertivores, and Combined Carnivores

**DOI:** 10.64898/2025.12.30.697105

**Authors:** Michael Tene, Kathleen Foley, Alexander Seaver, Wynn K Meyer

**Affiliations:** Lehigh University, Bethlehem, PA, United States; University of Iowa, Iowa City, IA, United States

**Keywords:** Diet, Molecular Evolution, Comparative Genomics, Mammals, Convergent Evolution, Genetics

## Abstract

Mammals have repeatedly evolved specialized diets, including a variety of predatory diets targeting different prey animals. Prior research has found differences in positive selection, gene family evolution, and gene functional loss across diets, but has primarily classified mammals into the three categories of herbivory, carnivory, and omnivory. Here we divide “carnivores” into vertivores and invertivores, due to the differences in nutrient composition of those food sources. We find significant differences in molecular evolutionary rates of multiple genes and gene sets between vertivores and invertivores, with vertivores having lower evolutionary rates of immune related genes, and invertivores having lower evolutionary rates of DNA repair genes and olfactory genes. By comparing evolutionary rates of vertivores, invertivores, and the combination of all carnivores against herbivores, we find that the method of classifying predatory species influences findings of differences between predators and herbivores. Our results in the combined carnivore-herbivore comparison are more similar to those of the invertivore-herbivore comparison than to those of the vertivore-herbivore comparison, which suggests that prior studies on carnivory may have been detecting patterns of molecular evolution related to invertivory. Lineages across all predatory diets have lower evolutionary rates of lipid metabolism genes relative to herbivores, though the specific genes and pathways showing these signatures, particularly those related to steroid hormone metabolism, differ between predatory diets. Taken together, these analyses of molecular evolutionary rates across nuanced predatory diets provide novel insights into the evolution of vertebrate- and invertebrate-based diets, as well as carnivory as a whole.

**Significance Statement:** Several studies have used genome-wide gene sequences from mammals with different diets to identify genes with patterns of natural selection related to diet. However, these studies were limited in part due to the common choice to combine invertebrate-based (e.g., insects, shellfish) and vertebrate-based (e.g., fish, birds, other mammals) diets in their analyses. By separating these diets in our study, we identify genes that experienced selection related to these different food sources across mammals, and provide clarification of patterns found by previous studies.

## Introduction

Diet has been a major selective pressure in vertebrate evolution and has led to numerous adaptive radiations (Grant and Grant 2006; Grundler and Rabosky 2021; Ocampo et al. 2022; Pandelis et al. 2023), including several in mammals (Monteiro and Nogueira 2011; Slater and Friscia 2019; Coombs et al. 2022). As mammalian lineages diverged from the ancestral insectivorous diet, they evolved many physiological, morphological, and behavioral adaptations associated with new diets (Casotti et al. 2006; Iason and Villalba 2006; Karasov et al. 2011; Emerling et al. 2018). Thus, diet influences selection on many aspects of mammalian ecology. Additionally, several diets have evolved multiple times in the mammalian phylogeny, including transitions back to insectivory (Wu 2022). This combination of many related biological functions and convergent evolution makes diet a strong candidate for using shared molecular patterns to connect evolution of genes with a phenotype using convergent evolution, an approach termed ‘PhyloG2P’ (Smith et al. 2020; Macdonald et al. 2025).

Several comparative genomics studies have assessed signatures of selection on a single branch of the mammalian phylogeny that represents an inferred dietary transition (Kim et al. 2016; Wang et al. 2016). This allows these studies to find genes with evidence of adaptation to the diet of interest. However, because these studies focus on members of one individual clade as representatives of their dietary strategy, it is unclear if the molecular evidence of adaptation is related to diet or other clade-specific traits. To address this limitation, other studies have examined diet-associated molecular evolutionary patterns across species from different clades that have independently evolved the same diet. With the inclusion of multiple independent clades, the number of shared features other than diet among foreground species is reduced, and thus the likelihood that the results are driven by a trait other than diet is also reduced. Two recent studies implemented genome-wide tests for differences in relative evolutionary rates across mammalian diets, considering diet as either a continuous or a categorical trait (Pollard et al. 2024; Redlich et al. 2024). Another examined convergent gene losses separately in herbivores and carnivores using a genome-wide approach (Hecker et al. 2019). However, a major limitation across these works is the use of a single diet called “carnivory” to describe all species that have predatory diets.

This simplification of carnivory highlights one of the major stumbling blocks in studies of diet evolution, defining diet categories. A common method of dividing diet is by trophic level, i.e., into herbivores, omnivores, and carnivores (Price et al. 2012; Price and Hopkins 2015; Famoso et al. 2018). However, previous work has demonstrated the importance of dietary variation not captured by trophic level and the limits of this simplification (Cantalapiedra et al. 2014; Pineda-Munoz and Alroy 2014; Reuter et al. 2023). Prior studies have taken alternative approaches to avoid the limitations of trophic level definitions through examination of carnivory as a continuous trait (Pollard et al. 2024) and use of more specific discrete diet categories or trophic guilds (Cantalapiedra et al. 2014; Meloro et al. 2015; Hargreaves et al. 2017; Miller and Pittman 2021; Reuter et al. 2023). These nuanced approaches have been key to refining our understanding of the evolution of mammalian diets, including through studies of niche definition, fossil analysis, and morphological convergence (Kelley and Motani 2015; Machovsky-Capuska et al. 2016; Chen et al. 2019). This suggests that a nuanced definition of diet is also important for studies of molecular evolution.

Although both vertebrate- and invertebrate-consumption have been considered “carnivory” in many prior works (Price et al. 2012; Li and Zhang 2014; Price and Hopkins 2015; Hecker et al. 2019; Román-Palacios et al. 2019; Mu et al. 2021; Stephens et al. 2022; Wagner et al. 2022; Wu 2022; Barts et al. 2025; Wilhoit et al. 2025), vertebrate and invertebrate food sources have distinct nutrient compositions, and their predators often use distinct prey acquisition strategies (Lintulaakso et al. 2023; Mayani-Parás et al. 2023). For instance, the protein content of vertebrate prey ranges from 40% to 80%, while invertebrate protein content ranges from 19% to 40% (Dierenfeld et al. 2002; Eisert 2011; Coogan et al. 2014; Senti and Gifford 2024). For these reasons, distinguishing between invertivorous and vertivorous animals is common in studies examining dietary diversity across species (Dierenfeld et al. 2002; Lintulaakso et al. 2023; Mayani-Parás et al. 2023; Senti and Gifford 2024). However, this distinction has so far been absent in prior work examining molecular evolution associated with diet, largely due to the limited number of species with genomic data available for analysis. Recent advances in tools designed for analysis of categorical traits (Redlich et al. 2024) and the availability of a larger dataset (Christmas et al. 2023) have made molecular evolution analysis with an invertivore-vertivore distinction feasible.

One of the challenges when working with differing diet definitions is a lack of standardization in terminology. While some comparative genomics studies refer to secondary consumers using the term ‘carnivore’ (e.g. (Hecker et al. 2019)), other works have used ‘carnivore’ to refer strictly to vertivorous species (e.g. Kim et al. 2016). To avoid ambiguity, in this paper we will use specific terminology described as follows. First, we use “invertivore” rather than the more common “insectivore” to describe predatory species that consume invertebrates, as our data includes species that eat non-insect invertebrates, such as krill-consuming cetaceans. Second, we use “vertivore” to refer to species that consume vertebrates. Third, we use the term “combined carnivore” to refer to the set of species that includes invertivores, vertivores, and species that eat a combination of vertebrates and invertebrates. “Combined carnivory” is equivalent to the trophic levels of secondary (and higher) consumers and scavengers. We additionally refer to “invertivore” and “vertivore” as “nuanced predatory diets,” in contrast to the combined carnivore diet. Finally, “invertivore”, “vertivore”, and “combined carnivore” are referred to collectively as “predatory diets”.

The objectives of this study are twofold. First, we examine molecular evolution using the nuanced distinction between vertivory and invertivory to identify genes with differences in relative evolutionary rates—implying different functional importance—between species with these predatory diets. Second, we compare which genes show differences in relative evolutionary rates in pairwise comparisons between herbivory and (i) invertivory, (ii) vertivory, and (iii) a single “combined carnivory” diet. This allows us to assess the consequences of the commonly used trophic level “carnivory” classification for analyses of molecular evolutionary rates associated with diet. We find substantial differences among the predatory diets in the analyses, with vertivory and invertivory showing distinct patterns, and combined carnivory resembling invertivory more than vertivory.

## Results

### Ancestral reconstruction of mammalian diet suggests multiple convergent dietary transitions

Because our molecular evolutionary analyses require inference of phenotypes for internal nodes of the phylogeny, we inferred ancestral diets using information for extant species. We determined the diet of 405 extant mammalian species using data from the EltonTraits dataset (Wilman et al. 2014), and classified these species into the categories: invertivore (68 species), vertivore (57 species), herbivore (196 species), and omnivore (84 species) (Supplementary Table 1). We used those diet classifications, along with known diets from the fossil record, to perform a maximum likelihood ancestral reconstruction of the diet for the internal nodes of the mammal phylogeny (Gill et al. 2014; Rothman et al. 2014; Emerling et al. 2018; Amador and Giannini 2021; Sadier et al. 2021). We then filtered our dataset to a smaller dataset of 193 extant species containing 67 herbivores, 42 invertivores, 57 omnivores, and 27 vertivores. This filtering allowed us to reduce the impact of over-represented clades on the analyses and remove short branches within clades sharing the same phenotype that provide limited added power in branch-based tests. This filtered dataset resulted in an inferred set of 137 herbivorous lineages (branches), 89 invertivorous lineages, 105 omnivorous lineages, and 52 vertivorous lineages within the tree. To assess whether results were robust to species selection during filtering, we performed the analysis with 100 alternatively filtered species sets (see Methods and Supplementary Text). The results reported here are significant (P ≤ 0.05 after Benjamini-Hochberg adjustment) in the primary analysis and at least 50% of the alternative species sets unless explicitly described as not robust (NR).

Mapping diet onto the phylogenetic tree reveals multiple convergent transitions among the four diets (Figure 1, Supplementary Data 5). While we observe an omnivorous intermediate in some clades with paths between predatory diets and herbivory (e.g., Caniformia, Xenarthra), for others we do not observe an omnivorous intermediate (e.g. Cetartiodactyla, Afrotheria). The lack of omnivorous intermediates may either represent direct transitions between the diets or be an artifact of each branch representing only the single inferred or observed phenotype of the downstream node or tip. Given that models of mammalian dietary evolution generally support omnivory as a near universal intermediate between herbivory and predatory diets (Price et al. 2012; Reuter et al. 2023), it is likely that these clades had an omnivorous ancestor but do not have any extant omnivorous lineages in our dataset. Our ancestral state reconstruction is based on phenotypes of extant species, along with select ancestral nodes with phenotypes shared by extant descendants, and thus any such clade would appear in the inferred “phenotype tree” (shown in Figure 1) to have experienced a direct transition between herbivory and a predatory diet. As the potential omnivorous intermediate would be present on only one branch in each such case, these rare potential artifacts are unlikely to affect downstream analyses which include many branches.

**Figure 1:**
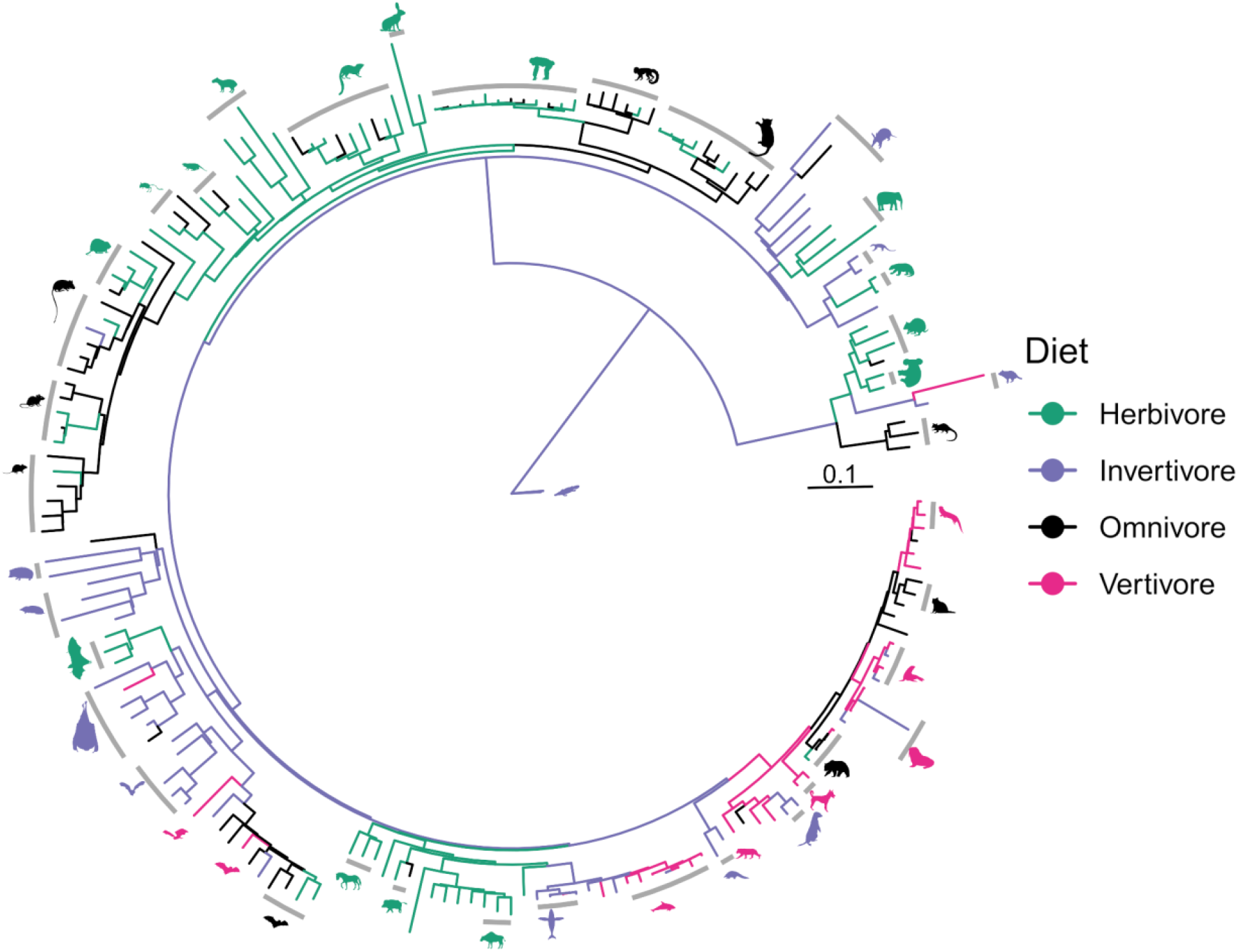
Inferred evolution of diet across the ancestors of 193 mammalian species. Phylogenetic tree of 193 mammalian species with branch lengths based on a subset of 16,209 filtered Zoonomia protein coding alignments (Wirthlin et al. 2019; Christmas et al. 2023; Kirilenko et al. 2023), with diets based on diet percentage data from EltonTraits (Wilman et al. 2014). Color indicates inferred or observed diet of the child node or tip of the branch. The scale bar represents a rate of 0.1 substitutions per site. Silhouettes from PhyloPic (<u>attribution</u>). See Table S1 for all species names and Data S5 for the tree.

### Vertivores and invertivores have differences in relative evolutionary rates across multiple biological functions

In order to identify genes undergoing changes in selective pressure associated with diet, we implemented a genome-wide test for diet-associated variation in relative evolutionary rates using RERconverge (Partha et al. 2019; Kowalczyk et al. 2020; Redlich et al. 2024), with the reconstructed diet evolution described above. We performed this analysis twice: the three diet analysis, where the diets of all 69 species with predatory diets were grouped as ‘combined carnivory’; and the four diet analysis, where vertivory (27 species) and invertivory (42 species) were considered as distinct diets (Figure 5 within Materials and Methods). Significant results from these analyses represent strong candidates for genes (Supplementary Data 1) and K-R categories (Supplementary Data 2, 3, 4) repeatedly responding to dietary selection across mammals.

To assess our hypothesis that vertebrate and invertebrate diets impose different selective pressures on a subset of genes, we directly compared the relative evolutionary rates of genes between vertivorous and invertivorous lineages. We found 21 genes and 13 KEGG-Reactome categories (K-R categories) with different relative evolutionary rates between lineages with the two diets (FDR = 0.05) (Supplementary Data 1, Supplementary Table 5). To aid in interpretation, we grouped the K-R categories into clusters based on shared genes (Figure 2). This produced 2 clusters that have lower relative evolutionary rates in vertivores than in invertivores, and 2 clusters that have lower relative evolutionary rates in invertivores than in vertivores. These clusters represent overlapping sets of genes with evidence of stronger purifying selection, and likely greater functional importance, in vertivores than in invertivores and vice versa. A complete representation of all results for tests of enrichment of K-R categories can be found in Supplementary Data 2.

**Figure 2:**
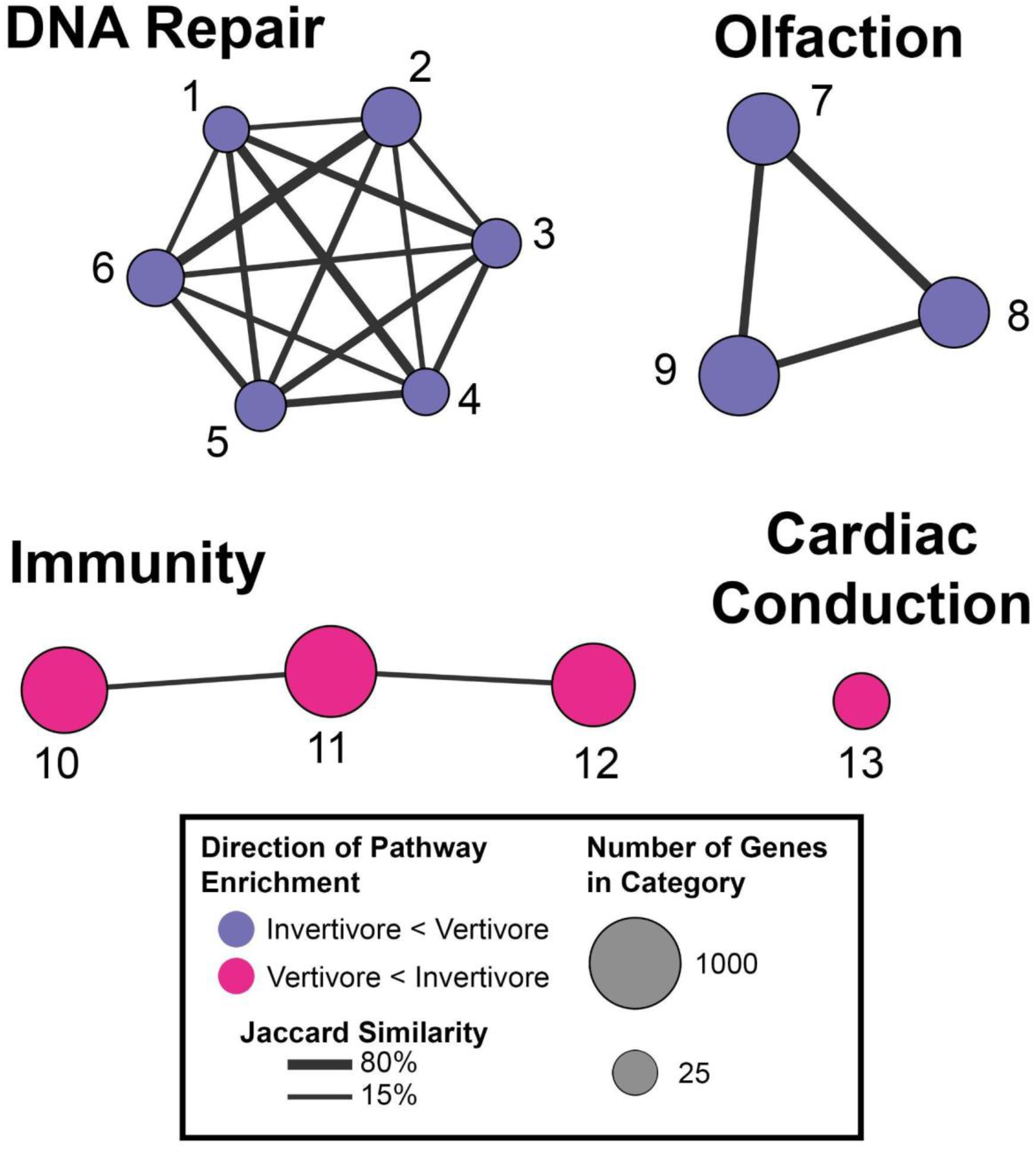
Clusters of K-R Categories evolving at different rates between vertivores and invertivores. Network graph of K-R categories enriched for genes with differing relative evolutionary rates (RER) between vertivores and invertivores at FDR = 0.05 and robust to species set selection. Clusters were generated using the igraph R package (Csardi and Nepusz 2005). Each node represents a K-R category. Nodes are connected by lines with thickness representing the percentage of genes shared between the nodes. Color indicates the direction of differences in RERs — if the category was enriched for genes evolving at a slower rate in vertivores (magenta) or invertivores (blue). Individual category names can be found in Supplementary Table 5.

Several K-R categories that are enriched for genes with lower relative evolutionary rates in vertivores than in invertivores are involved in or related to immunity, such as *Reactome Adaptive Immune System* (Figure 2 category 10, Mann-Whitney U AUC = 0.045, FDR-adjusted P = 2.6 × 10^−4^), *Infectious Disease* (Figure 2 category 11, AUC = 0.046, P = 6.1 × 10^−5^), and *Cytokine Signaling in Immune System* (Figure 2 category 12, AUC = 0.050, P = 7.6 × 10^−5^) (Supplementary Table 5, Supplementary Data 2). This strongly suggests that purifying selection to maintain immune response is stronger in vertivores than in invertivores, potentially due to differences in pathogen exposure. In the opposite direction, categories with functions related to DNA repair (6 categories, AUC = 0.09 - 0.24, P < 2.5 × 10^−4^), and olfaction (*Reactome Olfactory Signaling Pathway*, AUC = 0.14, P = 3.1 × 10^−6^; *KEGG Olfactory Transduction*, AUC = 0.10, P = 1.0 × 10^−4^; and *Sensory Perception*, AUC = 0.068, P = 9.1 × 10^−5^) are enriched for genes evolving slower in invertivores than in vertivores (Figure 2, Supplementary Data 2). These clusters are distinct, without shared genes or categories driving the signals.

A prior study working with an overlapping subset of species identified differences in active transposon content in the genomes of combined carnivores compared with those of herbivores or omnivores (Osmanski et al. 2023). Since recurrent transposon insertion could create selection on DNA repair genes (Munoz-Lopez and Garcia-Perez 2012), we examined whether this could explain the variation in relative evolutionary rates of these genes across diets in our dataset. When we re-examined transposon content with invertivores and vertivores as separate diets, we found significant variation across diets (*χ*^2^ = 32.7, P = 3.8 × 10^−7^), and in particular, greater transposon content in invertivorous than vertivorous lineages (Dunn’s Z = 4.1, P = 4.0 × 10^−5^) (Supplementary Table 9). We also found negative correlations between inferred mean transposon content and relative evolutionary rates of several DNA repair genes (e.g., *EME2, r* = −0.23, P = 0.044; *USP45*, *r* = −0.31, P = 4.7 × 10^−4^) (Supplementary Table 10, Supplementary Text). This suggests that differences in transposon content across diets could contribute to the observed slower evolution of DNA repair genes among invertivores in our dataset.

### Predatory diets show clear distinctions in comparisons of relative evolutionary rates with herbivory

Prior work has frequently compared molecular evolutionary patterns in species whose diets meet various definitions of carnivory with those of herbivorous species (Kim et al. 2016; Wang et al. 2016; Hecker et al. 2019; Pollard et al. 2024). Given the differences between vertivory and invertivory we observed here, we further examined how differences in the definition of carnivory can affect comparisons with herbivory. We compared the relative evolutionary rates of genes between herbivorous lineages and lineages from each predatory diet separately.

Among all three predatory diet-herbivore post hoc comparisons, combined carnivores had the highest number of genes with different relative evolutionary rates from herbivores (636, as compared to 405 for invertivore-herbivore and 139 for vertivore-herbivore), potentially due in part to the increased power from a larger sample size (141 combined carnivore branches from 89 invertivore and 52 vertivore branches). Increased power allows genes with a small difference in evolutionary rates in both the invertivore-herbivore and vertivore-herbivore analyses — for example due to a slight rate change within herbivores or rate changes of low magnitude in both invertivores and vertivores — to be detected by the combined analysis, but not the individual analyses. However, despite that increased power, the results from the combined carnivore-herbivore comparison did not contain all of the genes found in the comparisons of relative evolutionary rates between species with nuanced predatory diets and herbivores. This indicates that the nuanced diets demonstrate unique patterns of molecular evolution, while the combined diet remains useful for detecting shared patterns, especially subtle differences that require large sample sizes to detect. Lineages with nuanced predatory diets had many genes with significant differences in evolutionary rates relative to herbivores that were not significant in either the combined carnivore-herbivore analysis or the other nuanced predator diet-herbivore analyses (Figure 3).

**Figure 3:**
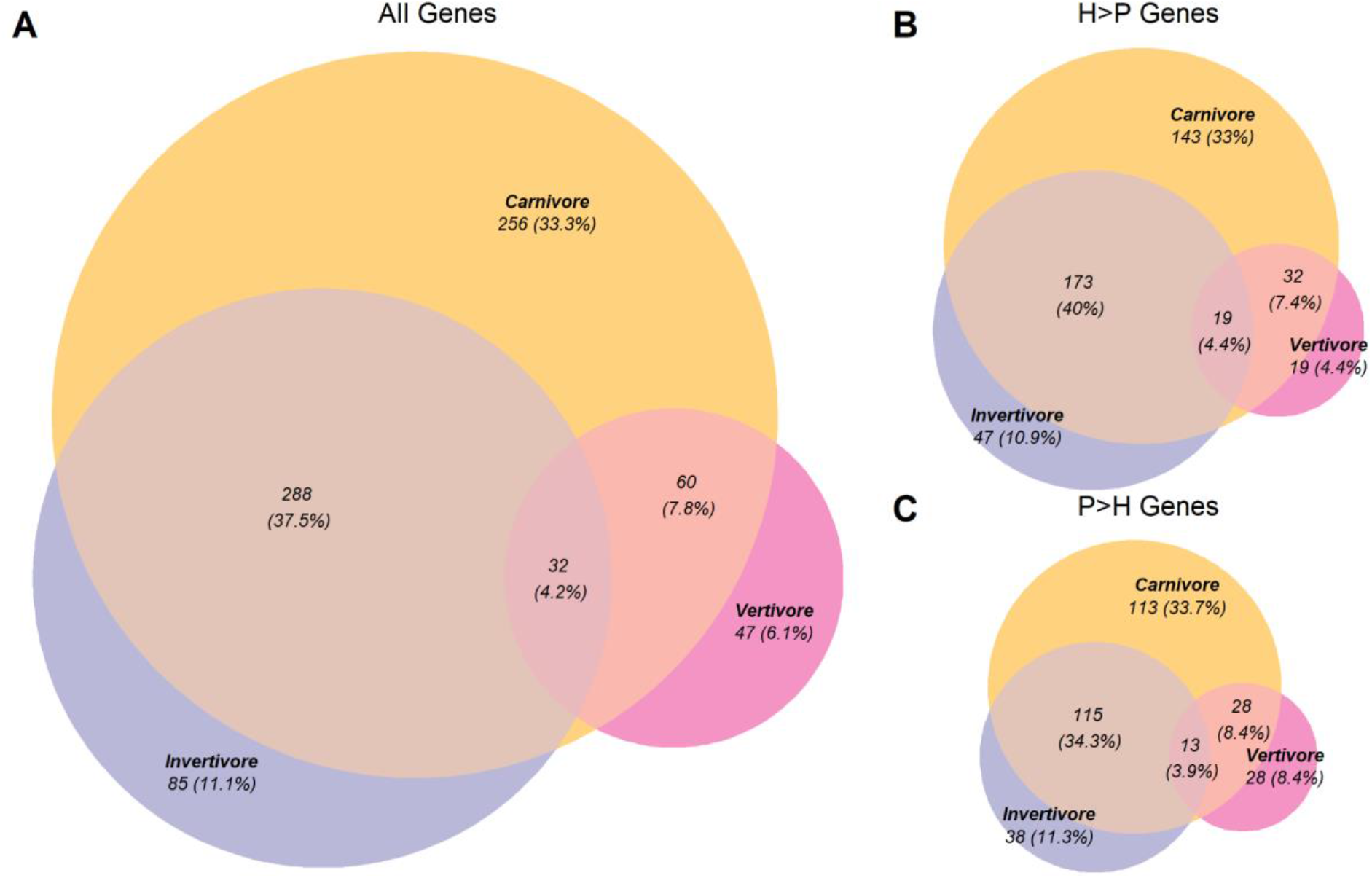
Limited overlap in evolutionary rate differences from herbivores between nuanced predatory diets. Representations of the overlap across analyses comparing relative evolutionary rates (RER) between herbivores and lineages with predatory diets. Genes included have FDR-adjusted P ≤ 0.05 and are robust to species set selection. Venn diagrams of (A) all genes with significantly different RER in each predatory diet than in herbivory, (B) genes with higher RER in herbivores than in species with predatory diets, and (C) genes with higher RER in species with predatory diets than in herbivores. Circle sizes represent the same scale throughout the figure. Note that percentages in the figure represent percentages of genes in each Venn diagram, while percentages in the text represent percentages of genes for individual diets.

Additionally, the overlap between the genes showing significant differences in the invertivore-herbivore and vertivore-herbivore comparisons was small, with only 23% of vertivore-herbivore results shared, and 7% for invertivore-herbivore. Overlap was higher but still limited at the level of K-R categories (60% of vertivore-herbivore, 2.5% of invertivore-herbivore) (Supplementary Figure 1, Supplementary Text). Taken together, these results suggest that combining invertivores and vertivores when comparing these species’ evolutionary rates to those of herbivores may obscure meaningful signals.

In these analyses, 320 (50%) of the 636 significant genes from the combined carnivore-herbivore analysis were shared with the invertivore-herbivore analysis, but only 92 (14%) were shared between the combined carnivore-herbivore analysis and the vertivore-herbivore analysis (Figure 3A). This pattern occurs both in genes evolving faster in species with predatory diets than in herbivores, as well as in genes evolving faster in herbivores than in species with predatory diets (Figure 3B, 3C). K-R categories also share a similar pattern of greater overlap between combined carnivore-herbivore and invertivore-herbivore results. This indicates that, within our dataset, combined carnivory reflects invertivory more than vertivory. This is likely due to the greater number of invertivores than vertivores in the mammalian species set we are using (89 invertivorous lineages, as compared to 52 vertivorous lineages, in the full tree). This greater abundance of invertivores is observed in Mammalia as whole, and thus represents a common pattern in pan-mammalian studies of diet (Pineda-Munoz and Alroy 2014; Reuter et al. 2023).

### Differences between vertivory and invertivory may influence prior work using combined carnivory

In previous work, Pollard et al. identified genes with variation in relative evolutionary rates associated with changes in a continuous definition of combined carnivorous diets in mammals, and they found six statistically significant genes: *ACADSB*, *CLDN16*, *CPB1*, *PNLIP*, *SLC13A2*, and *SLC14A2* (Pollard et al. 2024). Their analyses detected all of these genes as experiencing increasingly relaxed selective pressure with increasing herbivory, but they detected no signal of increasing purifying selection with increasing combined carnivory. Based on our findings of genes having differences in relative evolutionary rates relative to herbivores in either invertivores or vertivores but not combined carnivores, we re-examined the six significant genes from Pollard et al. (2024).

We found that all six genes were evolving significantly faster in herbivores than in combined carnivores; all of the genes were also evolving significantly faster in herbivores than invertivores (Table 1). One gene, *CLDN16*, was evolving faster in herbivores than in species with all other diets, including evolving faster in herbivores than omnivores (Dunn’s Z = 4.4, FDR-adjusted P = 4.8 ×10^−4^; Supplementary Data 1). These results are consistent with Pollard et al.’s findings of these genes experiencing increasingly relaxed purifying selection with increasing herbivory (Pollard et al. 2024). We note that not all of these results are robust to alternate species selection in our analyses; results for *CLDN16*, *CPB1*, *PNLIP*, and *SLC14A2* are robust to alternate species selection and permulations, but those for *ACADSB* and *SLC13A2* are not.

**Table 1.**
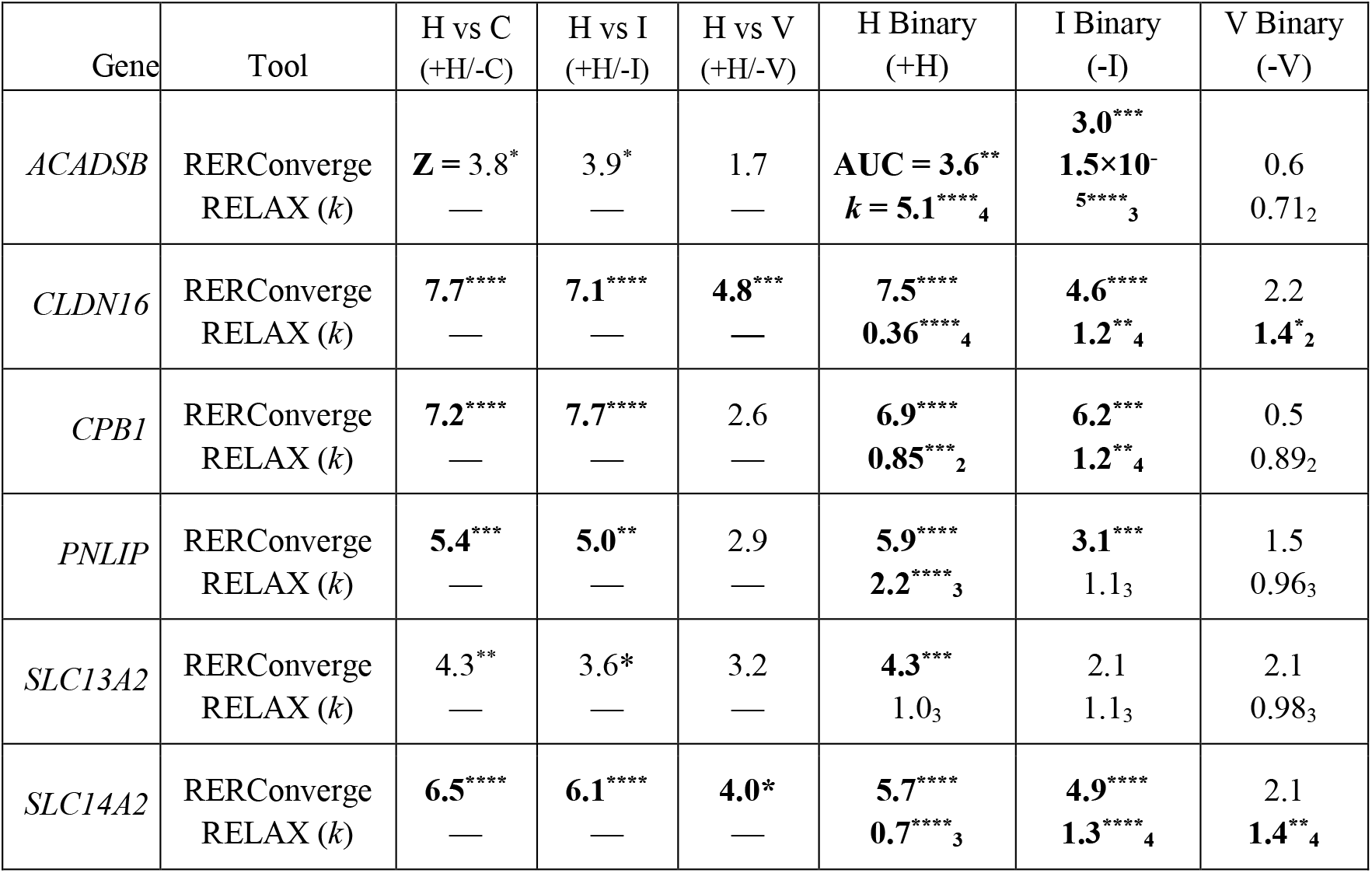
Nuanced diet analysis of genes with prior evidence of association between relative evolutionary rates and continuous combined carnivory. Genes identified by Pollard et al. as showing increasingly faster evolution with increasing herbivory, but not increasingly slower evolution with increasing combined carnivory (Pollard et al. 2024). RERconverge values are test statistics (Dunn’s Z for pairwise post hoc tests, Mann-Whitney U AUC for binary tests) representing the strength of association of changes in relative evolutionary rates (RER) between the listed diets. For RELAX intensification parameter *k*, subscript denotes optimal number of rate classes (see Methods). P-values listed are Benjamini-Hochberg adjusted P-values notated as P < 0.05 = *, P < 0.01 = **, P < 0.001 = ***, and P < 0.0001 = ****. Bolded values are significant and robust to alternate species selection. Signs and letters in parentheses indicate the direction of differences observed between categories; for example, +H/-C indicates that herbivorous branches had higher RER than carnivorous branches in the post hoc comparison, and -I indicates that invertivorous branches had lower RER than all other branches in the binary comparison. C: combined carnivore; H herbivore; I: invertivore; V: vertivore. A version of this table with the full values is included as Supplementary Table 8.

| Gene | Tool | H vs C<br>(+H/-C) | H vs I<br>(+H/-I) | H vs V<br>(+H/-V) | H Binary<br>(+H) | I Binary<br>(-I) | V Binary<br>(-V) |
|---|---|---|---|---|---|---|---|
| *ACADSB* | RERConverge<br>RELAX (*k*) | **Z =** 3.8<sup>\*</sup><br>— | 3.9<sup>*</sup><br>— | 1.7<br>— | **AUC = 3.6**<sup>\*\*</sup><br>***k* = 5.1**<sup>\*\*\*\*</sup><sub>4</sub> | **3.0**<sup>\*\*\*</sup><br>**1.5×10<sup>-5</sup>**<sup>\*\*\*\*</sup><sub>3</sub> | 0.6<br>0.71<sub>2</sub> |
| *CLDN16* | RERConverge<br>RELAX (*k*) | **7.7**<sup>\*\*\*\*</sup><br>— | **7.1**<sup>\*\*\*\*</sup><br>— | **4.8**<sup>\*\*\*</sup><br>**—** | **7.5**<sup>\*\*\*\*</sup><br>**0.36**<sup>\*\*\*\*</sup><sub>4</sub> | **4.6**<sup>\*\*\*\*</sup><br>**1.2**<sup>\*\*</sup><sub>4</sub> | 2.2<br>**1.4**<sup>\*</sup><sub>2</sub> |
| *CPB1* | RERConverge<br>RELAX (*k*) | **7.2**<sup>\*\*\*\*</sup><br>— | **7.7**<sup>\*\*\*\*</sup><br>— | 2.6<br>— | **6.9**<sup>\*\*\*\*</sup><br>**0.85**<sup>\*\*\*</sup><sub>2</sub> | **6.2**<sup>\*\*\*</sup><br>**1.2**<sup>\*\*</sup><sub>4</sub> | 0.5<br>0.89<sub>2</sub> |
| *PNLIP* | RERConverge<br>RELAX (*k*) | **5.4**<sup>\*\*\*</sup><br>— | **5.0**<sup>\*\*</sup><br>— | 2.9<br>— | **5.9**<sup>\*\*\*\*</sup><br>**2.2**<sup>\*\*\*\*</sup><sub>3</sub> | **3.1**<sup>\*\*\*</sup><br>1.1<sub>3</sub> | 1.5<br>0.96<sub>3</sub> |
| *SLC13A2* | RERConverge<br>RELAX (*k*) | 4.3<sup>**</sup><br>— | 3.6*<br>— | 3.2<br>— | **4.3**<sup>\*\*\*</sup><br>1.0<sub>3</sub> | 2.1<br>1.1<sub>3</sub> | 2.1<br>0.98<sub>3</sub> |
| *SLC14A2* | RERConverge<br>RELAX (*k*) | **6.5**<sup>\*\*\*\*</sup><br>— | **6.1**<sup>\*\*\*\*</sup><br>— | **4.0\***<br>— | **5.7**<sup>\*\*\*\*</sup><br>**0.7**<sup>\*\*\*\*</sup><sub>3</sub> | **4.9**<sup>\*\*\*\*</sup><br>**1.3**<sup>\*\*\*\*</sup><sub>4</sub> | 2.1<br>**1.4**<sup>\*\*</sup><sub>4</sub> |

In contrast to results from comparisons of herbivores with the other predatory diets, only two of these six genes (*CLDN16, SLC14A2)* had significantly faster evolution in herbivores than in vertivores (Z = 4.8, 4.0; P = 0.0035, 0.024), and this faster evolution was not robust to permulations.

To further examine how differences in evolutionary rates between vertivores and invertivores might influence these results, we performed a binary analysis comparing relative rates of these genes for branches with each individual diet against rates for all other branches. Supporting Pollard’s findings, all six genes were evolving significantly faster in herbivores than in all other species (Mann-Whitney U AUC = 3.7 - 8.0, FDR adjusted P ≤ 0.01) (Table 1, Supplementary Table 8). In addition, for five of the genes — *ACADSB*, *CLDN16*, *CPB1*, *PNLIP*, and *SLC14A2* — we also found significantly lower relative evolutionary rates in invertivores than in all other species in a binary analysis (AUC = 3.0, 4.6, 6.2, 3.1, 4.9; P = 0.030, 3.8 × 10^−4^, 1.8 × 10^−6^, 0.026, 1.5 × 10^−4^) (Table 1, Supplementary Table 8). This suggests an increase in the strength of purifying selection on these genes in invertivores, in addition to relaxation in herbivores. In binary analyses using HyPhy RELAX, we find that *CPB1* and *SLC14A2* both exhibit significantly relaxed selection in herbivores (*CPB1 k* = 0.85, P = 3.0 × 10^−4^, optimal rate classes = 2; *SLC14A2 k* = 0.73, P = 2.8 × 10^−15^, rate classes = 4), and *SLC14A2* exhibits significantly intensified purifying selection in invertivores (*k* = 1.3, P = 4.0 × 10^−7^, rate classes = 4), which aligns with this expectation (Table 1, Supplementary Table 2). Some of the RELAX results (for *ACADSB* in herbivores and invertivores, and *PNLIP* in herbivores) do not directly support the typical interpretation of the observed RERconverge patterns, i.e., that higher relative evolutionary rates indicate relaxed purifying selection. However, in each of these cases, the RELAX results appear to be driven by changes in the positive selection rate class (ω > 1) at a subset of sites. This is consistent with a different interpretation of the RERconverge results being driven by positive selection in herbivores on these genes.

Notably, these genes from the Pollard et al. study did not have markedly low relative evolutionary rates in vertivores, with none being significant in a binary analysis of vertivores versus all other species. This difference between the nuanced diets within the combined carnivory used by Pollard et. al. may explain their signal only being found as faster evolution with increasing herbivory, but not as slower evolution with increasing carnivory, as slower evolution was only occurring in a subset of their combined carnivores.

### Several metabolic pathways evolve at slower rates in predatory diets than herbivores, with diet-specific nuances

To assess the biological functions of genes with differences in evolutionary rates between predatory diets and herbivores, we identified KEGG-Reactome categories enriched for genes with differences in relative evolutionary rates between predatory diets and herbivores. We find several biological processes previously reported (Pollard et al. 2024) among those enriched for genes evolving more slowly in species with predatory diets than in herbivores, as expected, though we detect both previously reported and novel gene sets within the processes. These include lipid metabolism (e.g. *Reactome Metabolism of Lipids,* significant in all three predatory diet-herbivore post hoc comparisons: Mann-Whitney U AUC = 0.080 (combined carnivore-herbivore), 0.059 (invertivore-herbivore), 0.067 (vertivore-herbivore); FDR-adjusted P = 2.7 × 10^−9^, 3.8 × 10^−5^, 2.2 × 10^−5^); amino acid metabolism (e.g. *Reactome Metabolism of Amino Acids and Derivatives*, significant in combined carnivore-herbivore and invertivore-herbivore, AUC = 0.093, 0.090, 0.052, P = 3.0 × 10^−6^, 9.4 × 10^−6^, 0.06), biological oxidation (e.g. *Reactome Biological Oxidations* AUC = 0.13, 0.14, 0.09; P-value = 2.3 × 10^−8^, 2.5 × 10^−6^, 0.01 (NR)), and transmembrane transport (e.g. *Reactome Amino Acid Transport Across the Plasma Membrane*, significant in combined carnivore-herbivore, AUC = 0.17, 0.18, 0.16; P-value = 0.01, 0.009 (NR), 0.056) (Figure 4, Supplementary Table 6, Supplementary Data 2).

**Figure 4:**
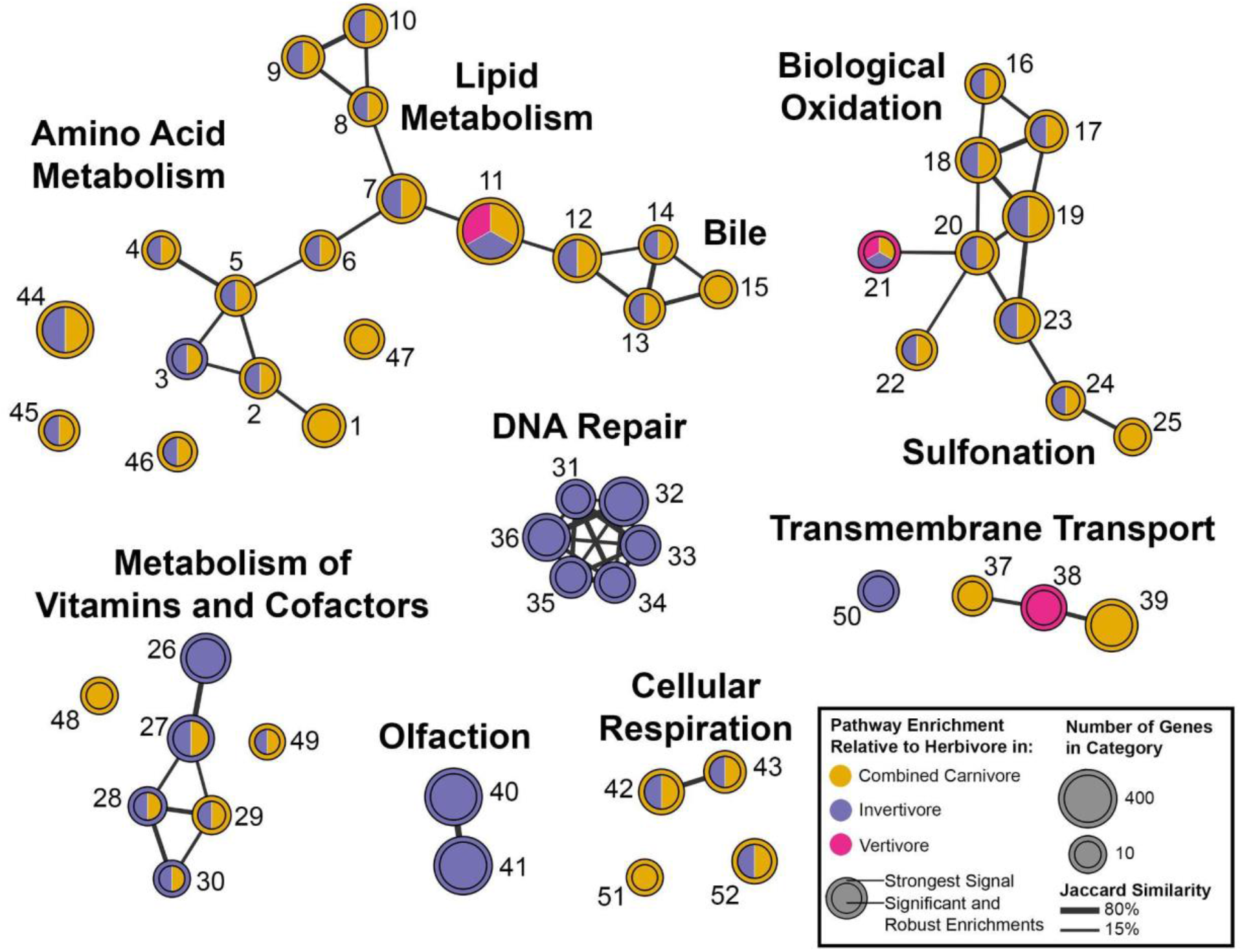
Multiple sets of pathways show lower relative evolutionary rates in predatory than herbivorous species. Network graph of K-R categories enriched for genes evolving more slowly in species with predatory diets than in herbivorous species, with at least one such enrichment significant at FDR = 0.05 and robust to species selection. Clusters were generated using the igraph R package (Csardi and Nepusz 2005) and labeled manually based on general function. Each node represents a K-R category; see Supplementary Table 6 for individual category names. Nodes are connected by lines with thickness representing the Jaccard similarity. Color within the circle indicates for which comparison(s) we observe significant enrichment (gold: combined carnivore-herbivore; blue: invertivore-herbivore; magenta: vertivore-invertivore). Circle outlines indicate the comparison with the strongest signal from the enrichment analysis.

We have also found novel pathways associated with dietary evolution, including cellular respiration (e.g. *Reactome Respiratory Electron Transport* AUC = 0.17, 0.15, 0.09; FDR-adjusted P = 5.3 × 10^−6^, 2.6 × 10^−4^, 0.1), peroxisome (e.g. *KEGG Peroxisome* AUC = 0.16, 0.16, 0.09; P = 3.4 × 10^−5^, 8.2 × 10^−5^, 0.13), and vitamin metabolism (e.g. *Reactome Metabolism of Water Soluble Vitamins and Cofactors* AUC = 0.11, 0.12, 0.04; P = 0.001, 7.5 × 10^−4^, 0.46) (Figure 4, Supplementary Figure 3, Supplementary Data 2, Supplementary Table 6). Separation of the nuanced predatory diets also allowed us to detect nuanced diet-specific categories within these clusters, such as the vertivore-specific *Reactome Transport of Inorganic Cations Anions* within transport (AUC = 0.08, 0.06, 0.1; P = 0.04 (NR), 0.13, 0.02) and the invertivore-specific *Reactome Metabolism of Vitamins and Cofactors* in vitamin metabolism (AUC = 0.08, 0.07, 0.03; P = 0.007 (NR), 0.019, 0.52).

While many broad patterns are shared across the predatory diets relative to herbivores, there is variation among the individual diets in the strength of enrichment for specific K-R categories. Some categories are clearly being driven more strongly by a single nuanced diet, such as *KEGG Steroid Hormone Biosynthesis*. While this category is enriched for genes evolving slower in predators than in herbivores in all three comparisons (FDR-adjusted P = 1.2 × 10^−4^, 1.2 × 10^−3^, 0.03), the herbivore-vertivore results show a stronger enrichment (Mann-Whitney U AUC = 0.28) than either the herbivore–combined-carnivore results (AUC = 0.22) or the herbivore-invertivore results (AUC = 0.16). An even more extreme case is *KEGG Tryptophan Metabolism*. It is enriched for genes evolving slower in both combined carnivores and invertivores than herbivores, but the invertivore-herbivore enrichment (AUC = 0.20, FDR-adjusted P = 8.9 × 10^−4^) is stronger than the combined carnivore-herbivore enrichment (AUC = 0.19, P = 6.5 × 10^−4^), and more than three times stronger than the vertivore-herbivore enrichment (AUC = 0.06, P = 0.57). This large enrichment difference suggests that genes involved in tryptophan metabolism may be evolving slower in only invertivores, not vertivores, and that this enrichment is significant in the combined carnivore comparison strictly due to the invertivores within combined carnivores. This large difference in enrichment is the case for all five categories relating to vitamin metabolism, suggesting that the entire cluster may be evolving slower in only invertivores.

In contrast, a large number of categories have a shared difference relative evolutionary rates between herbivores and all of the predatory diets. *Reactome Metabolism of Lipids*, for example, is strongly enriched in comparisons involving all of the predatory diets, with similar signal strength (combined carnivore-herbivore AUC = 0.081, P = 2.7 × 10^−9^, invertivore-herbivore AUC = 0.059, P = 3.8 × 10^−5^; vertivore-herbivore AUC = 0.067, P = 2.2 × 10^−5^). This pattern of shared enrichment includes some categories whose results are not significant in comparisons involving one or both of the nuanced diets. For example, *Reactome Recycling Of Bile Acids And Salts* is significant in only the combined carnivore-herbivore comparison (P = 0.01, 0.02 (NR), 0.08), but has similar enrichment in all three predatory diet comparisons (AUC = 0.23, 0.23, 0.21). This similar strength of enrichment in categories that are not significant in nuanced diet-herbivore analyses suggests a connection to the nuanced diet that the analysis is underpowered to detect. This pattern of similar signal strength occurs in the categories relating to amino acid metabolism (with the exception of tryptophan metabolism), as well as bile acids, the electron transport chain, biological oxidation, sulfur metabolism and cytochrome P450 (Supplementary Data 2, Supplementary Table 6). This is notable for the vertivore-herbivore comparison; it does not have any significantly enriched categories, but has consistently similar enrichment strengths, in these pathways.

*KEGG Steroid Hormone Biosynthesis* is the K-R category we found to be most strongly influenced by vertivory, with the highest AUC of all significant vertivore-herbivore enrichments (AUC = 0.28). To assess the potential biological relevance of this strong signal in vertivory, we examined the genes driving the enrichment for *Steroid Hormone Biosynthesis*. The enzymes driving the vertivore-herbivore enrichment in this category include two groups of genes: synthesis enzymes primarily expressed in the adrenal gland, and those primarily expressed in digestive tissues such as the liver, based on human RNA-seq datasets (Stelzer et al. 2016; GTEx 2025a; GTEx 2025b; GTEx 2025c; GTEx 2025d; GTEx 2025d).

Two of the genes expressed in the adrenal gland (*CYP17A1*, *CYP11A1*) are individually significant and robust, and another (*CYP21A2*) is not significant but highly ranked in the combined carnivore-herbivore (FDR-adjusted P = 1.5 ×10^−5^, 3.2 × 10^−4^, 0.006 (NR)) and invertivore-herbivore analyses (P =0.002, 0.006, 0.15). While these genes are not significant in the vertivore-herbivore analysis (P = 0.006 (NR), 0.07, 0.07), they have similarly high ranks in all three analyses (31, 107, and 293 in combined carnivore-herbivore; 162, 95, and 674 in invertivore-herbivore; 33, 267, and 274 in vertivore-herbivore) and thus contribute to the rank-based enrichment of steroid metabolism genes in all three analyses.

The other class of genes, hormone inactivation enzymes expressed in the digestive tract, were those with uniquely slow evolution in the vertivore-herbivore comparison. The top ranked gene in this category, *HSD11B1*, is individually significant in the post hoc vertivore-herbivore comparison (Dunn’s Z = 4.7, FDR-adjusted P = 0.004), and robust to alternate species selection but not to permulations (FDR-adjusted P = 0.42). This gene is not significant or highly ranked in the comparison of invertivores with herbivores (Z = 1.3, P = 1) and is not robust to alternate species selection in the combined carnivore-herbivore analysis (Z = 3.2; P = 0.047 (NR)), and thus this result is unique to the herbivore-vertivore comparison (Supplementary Data 2, Supplementary Table 6). *HSD11B1* (hydroxysteroid 11-Beta dehydrogenase) is involved in the conversion of the vertebrate stress hormone cortisol to the inactive metabolite cortisone, and is expressed in the liver and kidney (PubChem 2025). Two other digestive tract hormone processing enzymes (*HSD17B2*, *HSD17B3*) have larger differences in the vertivore-herbivore comparison (Z = 4.2, 3.7) than in the invertivore-herbivore comparison (Z = 2.3, 2.3), though the vertivore-herbivore comparison results are not individually significant (P = 0.018 (NR), 0.058). These two enzymes are both involved in androgen metabolism. *HSD17B2* is expressed in the liver and involved in testosterone inactivation; *HSD17B3* primarily performs testosterone activation in the testis, but also other androgen metabolism functions in the liver (GTEx Portal; GTEx Portal; PubChem; PubChem). The pattern of slower evolution in vertivores than herbivores of hormone inactivation and processing enzymes expressed in the digestive tract suggests that vertebrate-based diets may impose unique selective constraints related to processing ingested vertebrate hormones.

### Multiple signaling pathways and muscle contraction are evolving slower in herbivores than predatory species

We also find K-R categories that are enriched for genes evolving slower in herbivores than in species with predatory diets, suggesting that these processes are under stronger constraint in species with herbivorous diets (Supplementary Figure 3; Supplementary Table 7). A variety of K-R categories relating to different signaling pathways are significantly enriched for genes evolving slower in herbivores in the combined carnivore-herbivore and invertivore-herbivore analyses, such MAPK and WNT signaling, as well as other signaling pathways (*KEGG MAPK Signaling Pathway* AUC = 0.07, 0.07, 0.03; FDR-adjusted P-Value = 0.004, 0.008, 0.45; *Reactome Signaling By WNT* AUC = 0.10, 0.08, 0.08; P =1.0 × 10^−5^, 4.0 × 10^−4^, 0.016 (NR)) (Supplementary Data 2). Other members of these signaling pathways have been identified in previous comparative genomics analyses focused on diet (He et al. 2023; Redlich et al. 2024), with suggested explanations related to the pathways’ involvement in cell differentiation during digestive system development.

We also find K-R categories enriched for slower evolution in herbivores than in combined carnivores and invertivores in a variety of other processes, such as muscle contraction, senescence, cell junctions and adhesion, mRNA processing, and the neural system (*Reactome Muscle Contraction* AUC = 0.14, 0.13, 0.05; FDR-adjusted P = 2.3 × 10^−8^, 8.2 × 10^−7^, 0.18; *Reactome Cellular Senescence* AUC = 0.13, 0.13, 0.04; P = 1.2 × 10^−5^, 7.4 × 10^−6^, 0.45; *KEGG Tight Junction* AUC = 0.11, 0.08, 0.07; P = 0.001, 0.024, 0.18; *KEGG Spliceosome* AUC = 0.14, 0.16, 0.04; P = 0.001, 1.2 × 10^−4^, 0.63; *Reactome Neuronal System* AUC = 0.13, 0.13, 0.08; P = 2.2 × 10^−15^, 1.9 × 10^−12^, 1.3 × 10^−4^ (NR)). This implies that a wide range of biological processes — potentially including signaling, development, and neuron function — may experience stronger purifying selection in species with plant-based diets.

## Discussion

Here we find new insights into the evolution of diet by comparing the results of molecular evolutionary analyses using vertivores, invertivores, and a single category including all predatory diets. We find many differences in gene relative evolutionary rates between the predatory diets and in comparisons of each predatory diet with herbivory. Analyses that collapse predatory diets into a single category detect genes that neither of the nuanced diet analyses detect, while failing to detect many genes that are significant in nuanced comparisons of relative rates between invertivores and herbivores or between vertivores and herbivores. This suggests that the combined carnivore, nuanced invertivore, and nuanced vertivore diet definitions are all valuable, but not interchangeable. Given these findings, a similar nuanced approach may also be beneficial for analyses of herbivory. While our dataset was not well designed for such analyses, given the limitations of plant-based diet categorization in EltonTraits (Supplementary Table 1), future work to characterize the makeup of herbivorous diets in more detail may allow insights into the evolution of specialized herbivory.

Notable in light of the use of the term “carnivore” to refer to the combination of all predatory diets in other studies, we find that combined carnivory more closely reflects invertivory than vertivory in this large mammalian dataset; results for combined carnivores share more significant genes and K-R categories with invertivores than with vertivores in analyses relative to herbivores. We find several genes, K-R categories, and even clusters of K-R categories to be significant in the combined carnivore-herbivore analysis which are likely not associated with vertivory, and present only due to signal from invertivores. As this is due in part to the greater number of invertivores than vertivores in our dataset, we expect this pattern to be repeated in other pan-Mammalia analyses, because Mammalia as a whole has more invertivorous than vertivorous species (Pineda-Munoz and Alroy 2014; Reuter et al. 2023).

This greater similarity between combined carnivores and invertivores than vertivores is a concern because, in common vernacular, “carnivore” refers to the “consumption of flesh” as found in vertivory (Oxford Reference). In the most popular British, American, and learner’s English dictionaries, all ten examples included in the definition of carnivory were vertivores, not invertivores (Cambridge 2025 Oct 8; Merriam-Webster 2025 Oct 4; Oxford Reference). This strong link between “carnivore” and the consumption of vertebrates in the popular vernacular, as well as much of the scientific literature outside the niche of complex diet analyses, presents a serious communication challenge.

Despite reasonable approaches for avoiding confusion about combined carnivory, vertivory, and invertivory in evolutionary studies of dietary phenotypes, such as explicit descriptions of the definition of carnivory at the outset of a study, communication challenges persist. The first challenge is understanding when “carnivore” is being used to refer to “vertivore” in alignment with common vernacular, and when it is referring to the combination of predatory diets. When “carnivore” is being used to refer to a combination of predatory diets, it is additionally challenging for the reader to maintain the distinction between the combination of predatory diets and vertivores. Even if the distinction is explicitly highlighted by the author, the strong connection between “carnivore” and “vertivore” can persist for the audience, as new scientific knowledge often does not supplant earlier intuitions (Chuy et al. 2012; Shtulman and Valcarcel 2012; Stricker et al. 2021). This is especially true if the reader’s retention of the distinction is limited, such as during a cursory reading or when relying on long-term recall (Lustig et al. 2001).

We encourage researchers working on diets to consider these communication concerns in future work. The simplest solution may be to avoid using the term “carnivory” when describing diet, because of the ambiguous nature of the term. Possible alternatives include using ‘predator’ or ‘secondary consumer’ when referring to the combined diet, and vertivore and invertivore for the specific diets.

Making this distinction between nuanced predatory diets allowed us to identify genes and biological functions evolving at different rates in invertivores and vertivores. We find that genes and K-R categories related to immunity are evolving slower in vertivores than invertivores, and ones related to DNA repair and olfaction are evolving slower in invertivores than vertivores. Since we generally interpret lower relative evolutionary rates as the result of increased purifying selection, this implies that vertivores have experienced stronger purifying selection on immune-related genes than invertivores. This may be a result of vertivores experiencing increased exposure to pathogens through contact with their relatively closely related prey (Malmberg et al. 2021), an exposure not present in invertivores. The slower evolution of olfaction genes in invertivores may be related to the strong olfactory sense in some invertivores such as moles (Catania 2013; Catania 2020), though the particularly large gene family sizes and high turnover rates of olfactory genes pose challenges for interpretation (Kambere and Lane 2007; Hayden et al. 2010; Hughes et al. 2018). Future work incorporating models of gene family size change, as well as data for pseudogenes along with functional genes, might provide a clearer picture of the association between olfactory genes and dietary evolution in mammals.

Our comparisons of predatory diets with herbivores identified many pathways enriched for genes evolving slower in predatory species than in herbivores, including both previously reported pathways (e.g. amino acid metabolism, lipid metabolism, biological oxidation) and novel pathways (cellular respiration, peroxisome, vitamin metabolism, and DNA repair). For many of these K-R categories—such as the majority of those involved in lipid and amino acid metabolism—the species with predatory diets show similar differences in evolutionary rates from herbivores, suggesting a shared selective pressure across all predatory diets. Amino acids and lipids make up a larger proportion of predatory diets than herbivorous diets (Chizzolini et al. 1999; Karasov et al. 2011; Partha et al. 2019; Wu 2022). Thus, genes required for digestion of these compounds may be more relevant to fitness in predatory species than in herbivorous ones, due to differences in diet composition. Alternatively, these signals may be due to the different metabolic demands of predator and prey behavior and activity levels (de Matos Dias et al. 2018). Because many of the genes driving the enrichment of amino acid metabolism are directly involved in the catabolic pathways for these macromolecules (e.g. *AGXT*, *IDO, TAT, SDS*), this suggests that these signals represent the effects of diet composition. There may be selection on several biological processes underlying our observed lipid metabolism enrichment, since these gene sets contain some digestive or catabolic genes (*EHHADH*, *ABCB11, CUBN, HSD11B1*), but also genes relating to synthesis or broader biological functions (*CYP17A1*, *ALB, ACOT12*). The electron transport chain results are better explained by the differences in metabolism between species, as these genes (*ESCIT*, *NDUFB8*, *NDUFB5*) are involved in overall energy generation rather than the metabolism of specific macromolecules that differ between diets. Future work might explore the relative influences of diet composition and behavior on patterns of molecular evolution in metabolic pathways in mammals.

Notably, we find pathways and K-R categories that are likely to be driven exclusively by differences between invertivores and herbivores, despite being significant in the combined carnivore-herbivore analysis. These results highlight an interesting difference between vertivores and invertivores that would not be clear from combined carnivore analyses alone. The cluster of five categories related to vitamin metabolism shows this pattern of enrichment being driven specifically by evolutionary rate differences in invertivores (Supplementary Table 6, Supplementary Data 2). Several of the top genes driving enrichment of these pathways are involved in the transport of vitamins B6, B12, C, and D, or are metabolic enzymes that require these vitamins as cofactors (Watkins and Rosenblatt 2011). Specifically, several genes driving the enrichment of these pathways are essential in the vitamin B12-dependent conversion of methylmalonyl-CoA to succinyl-CoA (e.g., *ABCD4*, *AMN*, *CUBN*, *MMUT*, and *TCN1*) (Fyfe et al. 2004; Watkins and Rosenblatt 2011; Coelho et al. 2012). One potential reason for the observation of low relative evolutionary rates in invertivores for these genes is selection to prevent the buildup of methylmalonic acid. Mutations in a subset of these genes in humans have been linked to methylmalonic acidemia, a metabolic disorder characterized by the failure to properly break down protein and lipids in blood and tissues, resulting in a toxic accumulation of methylmalonic acid (Baumgartner et al. 2014). Invertebrates, such as marine mollusks, are known to contain large amounts of polyunsaturated fatty acids (Zhukova 2019), which can break down into methylmalonic acid precursors. This suggests the hypothesis that selection for efficient conversion of methylmalonyl-CoA to succinyl-CoA is stronger in invertivores than in other species, due to the greater proportion of methylmalonyl-CoA precursors in invertivorous diets. This could result in the reduced evolutionary rates we detect in invertivores relative to other species.

In the opposite direction, we find a variety of signaling pathways, as well as muscle contraction and mRNA processing, to be evolving slower in herbivores than in species with predatory diets. The slower evolution of muscle contraction genes in herbivores could represent increased purifying selection on muscle function in herbivores. A possible explanation for this change in selective pressure is differences in the gut between herbivores and other species (Duque-Correa et al. 2021); however, *Reactome Striated Muscle Contraction* and *Cardiac Conduction* are both significant in the combined carnivore-herbivore and invertivore-herbivore analyses, while *Reactome Smooth Muscle Contraction*, which is involved in gut function (Bitar 2003), is not significant. Alternatively, this decreased evolutionary rate in herbivores relative to predators may represent increased sequence change in predators due to positive selection. Positive selection on muscle genes has been reported in felids relative to herbivores, attributed to agile hunting behaviors (Kim et al. 2016). However, this positive selection was not detected in broader analyses of combined carnivory including predators with more diverse hunting strategies, a species set more comparable to ours (Pollard et al. 2024). Some of the signaling pathways we detected have been implicated in prior work on diet, such as MAPK and WNT signaling being involved in digestive tract development (Redlich et al. 2024). Another significant signaling type, GABA signaling, may have a more direct connection with diet as it shares genes with *Integration of Energy Metabolism* and *Regulation of insulin Secretion*, two other pathways evolving slower in herbivores than predators. The nervous system development and neural signaling pathways, however, lack a clear connection with diet. They may be being driven by appetite signaling, as suggested by findings from convergent loss of function studies (Hecker et al. 2019), or by differences in predator-prey behavior.

Another result without an immediate interpretation as a response to changes in diet is the lower evolutionary rates of DNA repair genes in invertivores than in both herbivores and vertivores. At least one other study has found a link between DNA repair and diet, with DNA repair genes being expressed at lower levels in two bamboo-eating herbivores than in closely related vertivorous species and omnivorous outgroups (Ma et al. 2023). A different study, also working with a subset of Zoonomia species, found that the genomes of predatory species have greater young DNA transposon content than those of omnivores or herbivores; the authors suggested that this pattern could be due to horizontal transfer from transposon-rich animal diets (Osmanski et al. 2023). Theoretically, this could lead to intensification of purifying or positive selection on DNA repair genes to combat transposable element disruption of the genome (Novick et al. 2010; Munoz-Lopez and Garcia-Perez 2012). Our results suggest that among the species shared with this prior analysis, invertivores have higher levels of DNA transposons than either vertivores or herbivores; this pattern may have been masked in the earlier study by the use of a combined carnivore category. We also see a negative correlation between relative evolutionary rates of DNA repair genes and young DNA transposon content within our dataset (Table S10). This suggests that invertivores may experience increased purifying selection on DNA repair genes due to dietary transposon content.

One notable difference we found between vertivores and invertivores relative to herbivores is in the genes responsible for the inactivation of vertebrate hormones in the digestive tract. The inactivated hormones – cortisol, estrogen, and testosterone – are all vertebrate-exclusive hormones (Jones and Reynolds 1980; Sapolsky et al. 2000; Hillier 2017; Nijhout and Laub 2018). The increased importance of hormone degrading enzymes in vertivores relative to invertivores could be a result of either the ingestion of the hormones, or differences in behavior and cognition between predator and prey animals. The first interpretation (that the signal is related to hormone ingestion) is supported by the observed patterns for genes expressed in different tissues. Specifically, relative evolutionary rates for the hormone synthesis enzymes expressed in the adrenal glands differ between herbivores and all predatory species, but rates for the inactivation enzymes expressed in the liver and digestive tract differ only between herbivores and vertivores. The fact that vertivores ingest vertebrate hormones while invertivores do not may explain the observed differences in digestive tract-expressed inactivation enzymes. Differences in behavior between predator and prey animals, however, may explain the shared difference in relative evolutionary rates of the adrenal gland enzymes between species with predatory diets and herbivores.

Overall, our results illustrate striking differences in evolutionary rates in proteins involved in numerous biological processes across mammalian lineages with different diets. By partitioning one trophic level into distinct categories, we have clarified which dietary features more strongly predict relative evolutionary rates for known diet-associated pathways, such as various metabolic pathways. We have also uncovered additional processes that appear to be subject to differing strengths of purifying selection depending on diet, or a phenotype co-evolving with diet, including DNA repair and steroid hormone metabolism. These findings highlight the value of expanding phylogenetic genotype-to-phenotype approaches to larger numbers of phenotypes and species. As more genomic and phenotypic data become available, we anticipate that such approaches will yield continued insights into the evolution of diet, including the evolution of specialized herbivores and of other quantitative dietary components, as well as into other complex mammalian phenotypes.

## Materials and Methods

### Inferring evolution of diet on the mammalian phylogeny

To perform our analyses, we used filtered amino acid alignments of protein-coding genes from the Zoonomia project (Christmas et al. 2023; Kirilenko et al. 2023; Wirthlin et al. 2024), containing 454 species. We gathered diet data for the 405 of these species that were included in the EltonTraits database (Wilman et al. 2014), and we used this database to assign each species one of the following diet categories for each analysis: herbivore, omnivore, vertivore, or invertivore for the four diet analysis; and herbivore, omnivore, or combined carnivore or for the three diet analysis (Supplementary Table 1). Next, we used those diet classifications, along with known diets from the fossil record, to perform a maximum likelihood ancestral reconstruction of diet for the internal nodes of the mammal phylogeny containing all 405 extant species with assigned diet categories (Revell 2012; Louca and Doebeli 2018; Paradis and Schliep 2019; Redlich et al. 2024). We describe each step in this process in further detail in the sections below.

#### Tip species diet classification

Diet information was drawn from the EltonTraits database (Wilman et al. 2014). The diet categories in EltonTraits are Diet.Inv (invertebrates, including aquatic invertebrates such as krill), Diet.Vend (land endothermic vertebrates), Diet.Vect (land ectothermic vertebrates), Diet.Vfish (aquatic vertebrates), Diet.Vunk (uncategorized vertebrates), Diet.Scav (carrion and scavenged vertebrates), Diet.Fruit (fruits and drupes), Diet.Nect (nectar and other plant secretions), Diet.Seed (seeds and grains), and Diet.PlantO (other plant matter: grass, ground vegetation, seedlings, weeds, lichen, moss, small plants, reeds, cultivated crops, forbs, vegetables, fungi, roots, tubers, legumes, bulbs, leaves, above ground vegetation, twigs, bark, shrubs, herbs, shoots, aquatic vegetation, aquatic plants). We grouped these into Herbivorous Categories (Fruit, Net, Seed, PlantO), Vertivorous Categories (Vend, Vect, Vunk, Vfish, Scav), and Invertivorous Categories (Inv). These were used to assign diet categories (Figure 5).

**Figure 5:**
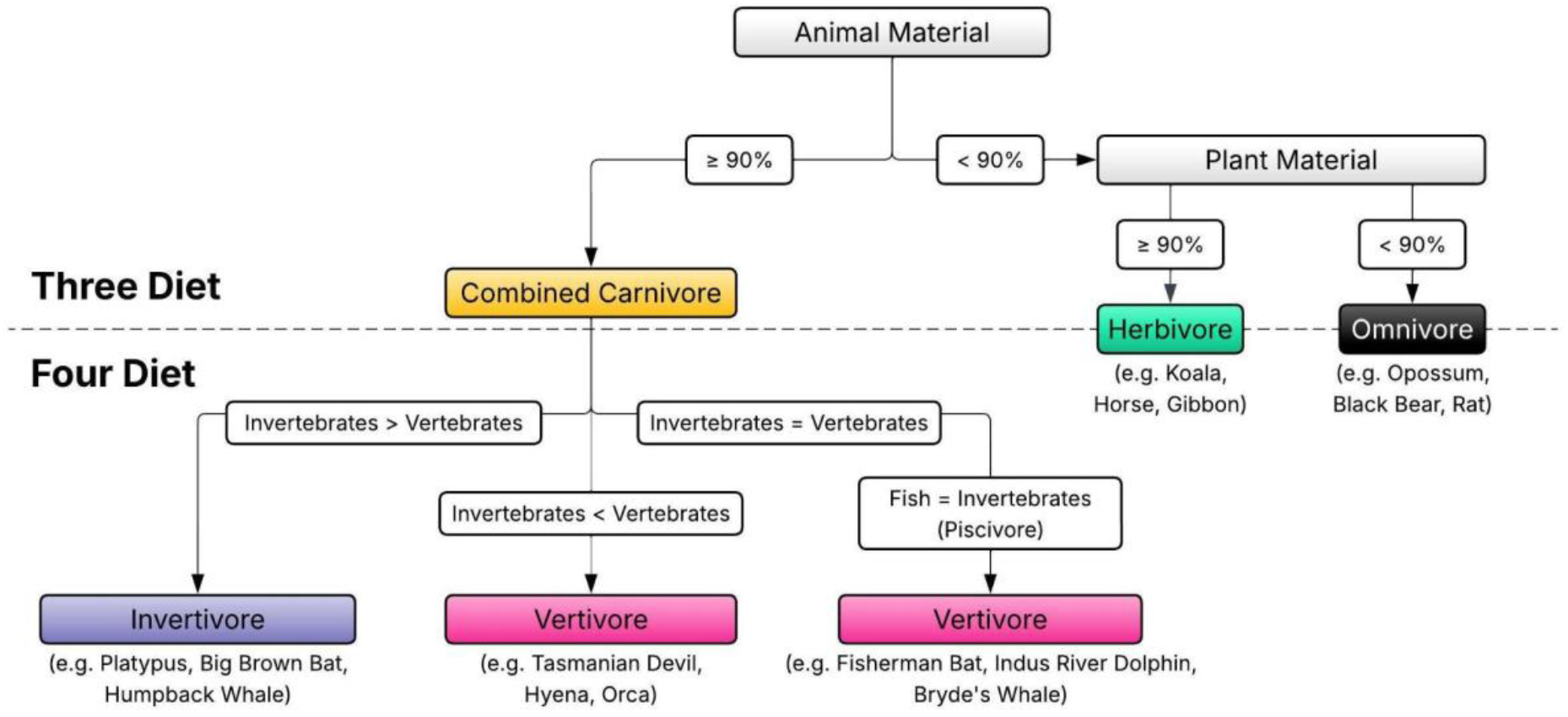
Schematic visualizing diet assignment process based on EltonTraits data. Animal material represents the combination of vertebrate and invertebrate material. The herbivore and omnivore classifications were used in both the three-diet and four-diet analyses. The combined carnivore classification was used in the three-diet analysis, and split into invertivore and vertivore in the four diet analysis. In our dataset, all of the species with equal vertebrate and invertebrate material were species with an equal amount of fish and invertebrate material, and thus considered piscivores.

In the three diet analysis, we assigned a diet using a 90% threshold of each of the above categories. We classified species with a diet composed of ≥90% herbivorous categories as herbivores. We classified species with a diet of ≥90% [vertivorous categories + invertivorous categories] as combined carnivores. All other species were classified as omnivores (Figure 5).

In the four diet analysis, we split combined carnivory into vertivores and invertivores. We classified the above defined combined carnivores that consume more than 50% vertivorous categories as vertivores, and those that consume more than 50% invertivorous categories as invertivores. Combined carnivores with exactly 50% vertivorous categories and 50% invertivorous categories consisted of exclusively species with a diet of 50% fish and 50% invertebrates, which are often considered piscivores (Kelley and Motani 2015; Aizpurua et al. 2021; Izadi et al. 2022; Das et al. 2025). Piscivory is often considered a subset of vertivory (Kissling et al. 2014); thus, we classified these species as vertivores, to maintain consistency with other work in the field. An alternative approach would be to exclude these species from the analysis, but we preserved them to maintain species set consistency with the three diet analysis. Instead, we performed a secondary alternative 4 diet analysis, in which combined carnivore species that did not pass the 90% threshold in either vertivorous or invertivorous categories were excluded from the analysis, and we assessed concordance of the results of this analysis with those of our main analysis (Supplementary Text).

We determined which threshold should be used for diet percentage cutoffs by identifying a threshold that was best able to predict other datasets. For this, we used the PanTHERIA database (Jones et al. 2009) as well as a database of classifications we created based on information drawn from Walker’s Mammals of the World (Nowak 1999); classifications from both of these sources can be found within Supplementary Table 3. Because PanTHERIA only included diet assignment at the trophic level, we compared our assignments for the three diet categories. We found that a ≥90% threshold was best able to predict diet assignments from the other datasets (246 of 314 match in PanTHERIA for the three diet categories, 179 of 216 match the database from Walker’s Mammals of the World). The results of our prediction accuracy analysis can be found in Supplementary Table 3.

#### Ancestral state reconstruction

We inferred ancestral diets for all internal nodes of a tree containing all 405 extant species with defined diets in our study (Supplementary Table 1). As input to ancestral state reconstruction, we used the master tree inferred from our protein-coding alignment data (see below), with a tree topology from the guide tree that was used for the MACSE alignments in the study first reporting the development of the TOGA software (Schliep 2011; Kirilenko et al. 2023; Hiller 2024). We initially inferred ancestral states for all internal nodes of this tree through a maximum likelihood ancestral trait reconstruction using a continuous time Markov model of evolution, implemented through the char2TreeCategorical function contained within RERConverge, with the default equal rates transition model (Revell 2012; Louca and Doebeli 2018; Paradis and Schliep 2019; Redlich et al. 2024). However, we checked these inferences against several ancestral diets strongly supported by the fossil record, and we found that this method was unable to correctly predict the known ancestral diets. This default ancestral state reconstruction method incorrectly inferred several ancestral nodes as being herbivores although the fossil records show them to be otherwise; specifically, the fossil record supports the ancestral mammal, ancestral marsupial, ancestral placental mammal, and the ancestral chiropteran as invertivores (Gill et al. 2014; Emerling et al. 2018; Amador and Giannini 2021; Sadier et al. 2021); and the ancestral primate as an omnivore (Rothman et al. 2014). As such, we implemented a method provided by Liam Revell to fix the values of those ancestral nodes to the diets of each that are best supported by the fossil record (Revell 2025). This was achieved by adding an additional tip at the ancestral node, creating a tritomy. This new tip had a branch length of zero, which caused the ancestral node to inherit the phenotype value of the new tip, set to the diet best supported by the fossil record. Ancestral reconstruction was then performed again using the char2TreeCategorical function within RERconverge, with the default equal rates transition model (Revell 2012; Louca and Doebeli 2018; Paradis and Schliep 2019; Redlich et al. 2024) with tritomies included in the tree, after which the tritomies were removed. This procedure successfully led to the inference of ancestral nodes best supported by the fossil record. The resulting categorical “phenotype tree” represents observed phenotypes of tip species and inferred phenotypes of ancestral nodes on upstream branches; branch values 1 - 4 represent phenotypes Herbivore, Invertivore, Omnivore, and Vertivore (Supplementary Data 5).

#### Binary “phenotype tree” creation

We created binary “phenotype trees,” or trees whose branch lengths represent background (1) or foreground (2) lineages, for use in the analysis of genes previously identified in Pollard et al. (Pollard et al. 2024), as well as for use in HyPhy RELAX analyses (see details in “Assessing relaxation or intensification of purifying selection on genes of interest” below). To generate these trees, we directly modified the branch values on the tree representing the evolution of the categorical phenotypes inferred using the ancestral state reconstruction method described above. For each of the four diet phenotypes, we created one binary phenotype tree with all branches of the categorical phenotype tree inferred as that phenotype set to foreground and all other branches set to background (Supplementary Data 5).

### Genetic Data and Gene Tree generation

#### Generation of topology-constrained gene trees

In order to estimate relative evolutionary rates for each gene that account for variation in rates across genes and branches, RERconverge requires that all gene trees have the same topology, representing the species tree (Kowalczyk et al. 2019). We generated a set of topology-constrained gene trees for our analyses using the TOGA-generated amino acid alignments for protein-coding genes available from the Zoonomia Consortium (Kirilenko et al. 2023), filtered as described in Wirthlin et al. (Christmas et al. 2023; Wirthlin et al. 2024) to remove low-quality sequences and alignments. We used each of these 16,209 filtered amino acid alignments as input to a phangorn-based maximum likelihood branch length inference method with LG as the substitution model, and with topology constrained by a user-provided guide tree (Le and Gascuel 2008; Schliep 2011). We provided the guide tree that was used for the MACSE alignments in the study first reporting the development of the TOGA software (Schliep 2011; Kirilenko et al. 2023; Hiller 2024). We then used the resulting topology-constrained gene trees to generate a “master tree;” the branch lengths of this master tree represented averages from all 10 gene trees that contained all of the 454 species in the filtered alignments. This master tree and the phenotype trees inferred as described in *Ancestral state reconstruction* above are available as Supplementary Data 5.

#### Species filtering

Of the 454 species in the filtered alignments, we were able to classify diet phenotypes for 405 species using the thresholding method with the EltonTraits data described above (Wilman et al. 2014). We filtered these 405 species to a smaller set of 193 species for use in subsequent analyses, choosing a subset of species to maintain a balance between large overall sample size and relatively even taxon sampling across mammalian clades. After all filtering steps described below, our species set contained 193 species provided to all RERConverge analyses using the useSpecies argument.

Our first filtering step was to remove any species present in too few protein-coding alignments, which may represent species with poor quality sequences in the alignments. This prevents later filtering steps from removing a species with data for a large number of genes in favor of a species with data for very few genes. In this step, we removed 5 species that were present in a number of alignments three standard deviations below the mean (<11,300 genes). We did not exclude the monotremes, despite them having sequences in the alignments of fewer than 11,300 genes, for three reasons. First, inclusion of monotremes allows us to increase the breadth of our analysis from a pan-therian analysis to a pan-mammalian analysis. Second, monotremes would not cause the aforementioned problem of removing a species with a large number of genes in favor of a species with fewer genes in a subsequent filtering step, as that only affects species with short branch lengths. Both monotremes are on terminal branches with lengths >0.01, and the subsequent branch length filters only apply to branches <0.01. Finally, monotreme exclusion from the alignments is less likely to represent poor quality sequences in monotremere than for other species, as monotremes are the most distantly related to the other species in the alignments and so are less likely to have orthologs identified.

Because the RERConverge method is based on testing for differences in evolutionary rates among branches in gene trees, short branches without sufficient time for the protein sequence to accumulate changes are unlikely to contribute substantially to the analyses and will often be masked by default when computing relative evolutionary rates (Partha et al. 2019). For this reason, we removed 143 short terminal branches (<0.01 amino acid substitutions/site in the master tree) that did not have a phenotype transition. We retained transitional branches in the main analysis in order to maximize the number of convergent phenotype transitions within our dataset; this allows the detection of repeated shifts in evolutionary rates across clades, with a tradeoff of some power loss due to short branch lengths. To select branches to be dropped, we iteratively assessed the phenotype state of the shortest terminal branch of the master tree, relative to the inferred state of its ancestral branch. If this terminal branch was involved in a phenotype transition, the branch was preserved. If not, the branch was dropped, and we again assessed whether the shortest terminal branch of the newly filtered tree was involved in a phenotype transition, excluding from this assessment any branches that had previously been preserved as transitional. We repeated this process until the filtered tree contained no terminal branches with a length of less than 0.01 not involved in a phenotype transition. Only species that remained in this pruned tree were included in the species set used in the main RERconverge analysis.

The intersection of species in the EltonTraits and Zoonomia datasets used as the input for our analysis lacks balanced taxon sampling. While 29 families contained invertivorous species, a single family, Vespertilionidae, included 13 of the 59 extant invertivores (22%) in the original data and 10 of the 49 extant invertivores (20%) after filtering for branch length. Five monophyletic clades that are exclusively herbivorous (families Bovidae, Cervidae, Cricetidae, and Pteropodidae, and infraorder Hystricognathi) included 91 of the 185 (49%) extant herbivorous species in our dataset before pruning for branch length (31 of the 95 extant species (32%) after pruning). Because these clades do not contain multiple diets, they introduce confounders to the analyses. Any non-diet features shared by such a clade would be present in only species with one diet, and this could cause genes whose relative evolutionary rates are associated with these clade-specific non-diet features to appear significant in the test for association between relative evolutionary rates and diet. An example is that 906 genes and 81 K-R categories have significantly different relative evolutionary rates between the 6 herbivorous rodents in Cricetidae and all other species, including other herbivores (Supplementary Data 6). These results represent differences in evolutionary rates that are specific to Cricetidae and are not necessarily related to herbivory. These Cricetid-specific rate differences could disproportionately influence the distribution of relative evolutionary rates across herbivorous lineages if a large percentage of herbivores were Cricetids. To minimize these issues, we reduced these large clades that were overrepresented within lineages consuming individual diets (Vespertilionidae in invertivores and Bovidae, Cervidae, Cricetidae, Hystricognathi, and Pteropodidae in herbivores) to the modal number of species per family of 3, selecting 3 species from each clade at random (Supplementary Table 1). This process was applied to the 6 overrepresented clades (families Bovidae, Cervidae, Cricetidae, Pteropodidae, and Vespertilionidae, and infraorder Hystricognathi), removing a total of 39 species.

### Analysis of relative evolutionary rate differences associated with diets

We performed the analysis of the association between gene relative evolutionary rates and diet using the categorical expansion of the RERConverge package (Redlich et al. 2024). The analyses were performed using new wrapper scripts for RERConverge functions, which can be found at https://github.com/wkmeyer-lab/RunRERPublication, along with the exact arguments provided to the wrapper functions included in the scripts.

We ran RERConverge using the categorical analysis method with the categorical and binary “phenotype trees” described above as input and default settings. Genes were included in a given analysis if the gene tree included at least 10 total branches, and at least 2 branches in each category. We performed the RERConverge analysis once using the 4 category diet classifications, once using the 3 category diet classifications, and once for each category as foreground in a binary analysis. Multiple hypothesis testing correction was performed within each analysis using a Benjamini-Hochberg adjustment (Benjamini and Hochberg 1995) with a FDR threshold of 0.05.

We performed gene set enrichment analysis using a Wilcoxon rank-sum (or Mann-Whitney U) enrichment test on gene set categories using the fastwilcoxGMTall function in RERConverge (Kowalczyk et al. 2019). We used the KEGG and Reactome gene sets from MSigDB 3.0 version 2023.1.Hs in this analysis (Kanehisa and Goto 2000; Liberzon et al. 2011; Kowalczyk et al. 2019; Milacic et al. 2024). KEGG and Reactome gene sets were identified by finding all gene sets using those prefixes from EnrichmentHsSymbolsFile.gmt from the GSEA database (Subramanian et al. 2005). We also performed this analysis on other gene sets (see Supplementary Results). Multiple hypothesis testing correction was performed separately for each set of gene sets using a Benjamini-Hochberg adjustment with a FDR threshold of 0.05.

To visualize gene set enrichment analysis results, we created clusters with the R packages ggraph and igraph (Pedersen and RStudio 2025; Csárdi et al. 2026) using the Fruchterman-Reingold force-directed algorithm (Fruchterman and Reingold 1991), filtered for edges containing at least 15% gene set overlap. We labeled the clusters manually based on the component categories, for use in visualization and interpretation.

### Assessing relaxation or intensification of purifying selection on genes of interest

In addition to RERConverge, we used the Hypothesis Testing using Phylogenies (HyPhy) tool RELAX (Pond et al. 2005; Wertheim et al. 2015) to detect relaxed or intensified selection for all genes present in Table 1 and select genes that drive enrichment of K-R categories from the post hoc comparisons in the RERconverge analyses. All genes analyzed with RELAX are listed in Supplementary Table 2 along with the RELAX results. For each gene, we used the codon alignment corresponding to the filtered amino acid alignment used for RERConverge as input to RELAX. Then, we used the binary “phenotype trees” described above as inputs to RELAX in order to set one of the four diet categories as test branches, with all other diets set as reference; we repeated this separately for each of the diets. We ran RELAX with the following parameters and settings: starting points set to 10, grid size set to 250; and we ran a two, three, and four rate class analysis for each diet. We report the parameter *k* and Benjamini-Hochberg FDR-adjusted p-value for the number of rate classes with the lowest AICc within the main text and Table 1. Additional information, including estimates of ω and the proportion of sites within each rate class for the lowest AICc result for each gene and test diet, can be found in Supplementary Table 2; all RELAX results are contained in Supplementary Data 7.

### Testing for robustness

To assess whether our categorical RERconverge results for the 3- and 4-diet analyses were robust to species selection resulting from the species filtering steps described above, we re-ran the analyses on 100 random sets of species matched for overall sample size and sample sizes within each diet category. These species sets were created to provide plausible alternate sets of species that could have been generated during the species filtering and tree pruning steps, without specific curation for transitional branches or overrepresented families. To create these alternate species sets, we selected random samples of 193 species from the 405 species in our unfiltered dataset, including in each alternate species set the same number of extant species from each category as were included in the main analysis (67 herbivores, 42 invertivores, 57 omnivores, and 27 vertivores).

A candidate set of species generated in this way was then used as an alternate species set if the number of short branches (branches with a length of <0.01, the branch length cutoff used in pruning) was less than 1.5 times the number of short branches in the main analysis. This increased number of allowed short branches in the alternates allows for a greater variety of possible species combinations within the alternate species sets and thus a wider variety of possible alternative pruning decisions. Supplementary Table 1 includes a column showing the number of alternate species sets (out of all 100 sets) in which each species is included.

When reporting significant and robust results in the main text, we include results that were significant at FDR-adjusted P ≤ 0.05 in at least half of the analyses of alternate species sets, in addition to the main species set analysis. We have included summary statistics (mean, median, standard deviation, and IQR) of the test statistic, P-value, and FDR-adjusted P-value for each gene and K-R category across the analyses of the 100 alternate species sets in our Supplementary Data (Supplementary Data 1, Supplementary Data 2).

#### Phylogenetic permulations

To account for the potential influence of the phylogenetic distribution of diet phenotypes on the null distribution of test statistics, we performed a phylogenetically restricted permutation strategy known as permulations (Kowalczyk et al. 2020; Saputra et al. 2021). This created “phenotype trees” with a number of branches per phenotype within 10% of those of the diets included in our analyses, but without any known biological factors in common across all branches assigned the same phenotype. Performing RERconverge analyses on these “null phenotype trees” can thus generate a null distribution of test statistics that includes the effects of realistic patterns of trait evolution on the tree. We performed 2,600,000 permulations of the phenotypes for species included in the main four-diet analysis, using the ER ancestral state reconstruction model and ‘auto’ root probability options in RERconverge version 0.1.0. We determined permulation-based P-values for each gene by computing

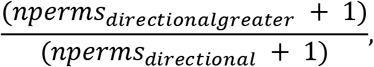

where *nperms_directional_* represents the number of permulations in which the direction of effect was the same as observed in the main analysis, where applicable; and *nperms_directionalgreater_* represents the subset of *nperms_directional_* in which the absolute value of the test statistic for a given gene (Kruskal-Wallis H or Mann-Whitney U AUC) in the permulation analysis was greater than or equal to that observed in the main analysis results for that gene. We implemented Benjamini-Hochberg corrections to adjust permulation-based P-values for the omnibus test and each pairwise post hoc analysis separately. Additionally, we assessed if the K-R categories that were significant in the main analysis and robust to species selection (FDR-adjusted P-value ≤ 0.05 in ≥50% alternate species sets) were also robust to permulations by performing the gene set enrichment using the permulation-based P-values. All of the robust K-R categories were significant in the permulations analysis, and unless explicitly stated otherwise, all individually significant genes discussed in the main text are significant in the relevant permulation analysis (FDR-adjusted permulation-based P-value ≤ 0.05). However, we report FDR-adjusted P-values from the main analyses, rather than permulation-based P-values, in the main text for consistency with the reported test statistics. We include permulation-based P-values in Supplementary Data 1 and 2.

#### Testing robustness to reconstruction method

To assess the robustness of our results to the choice of ancestral state reconstruction method, we compared our four-diet categorical RERconverge results with those obtained using a different inference method. The alternative inference method was the categorical RERconverge ancestral state inference method char2TreeCategorical (Revell 2012; Louca and Doebeli 2018; Paradis and Schliep 2019; Redlich et al. 2024) using an equal rates rate model and a tree with all equal branch lengths, chosen as it correctly predicted the ancestral diets best supported by the fossil record (insectivorous ancestor of Mammalia, Marsupialia, Placentalia, and Chiropreta; omnivorous ancestral Primate). This resulted in 11 branches of the “phenotype tree” (inferred ancestral nodes) with a diet that differed from that in the main analysis “phenotype tree” (Supplementary Data 5). The results from analyses using this alternative inference were largely concordant with those using the ancestral state reconstruction method we described for our main analysis. The Mann-Whitney U AUC and FDR-adjusted P-values of genes were strongly correlated between analyses using the two inference methods for all pairwise post hoc comparisons from the combined carnivore-herbivore (R^2^ = 0.99, 0.95), invertivore-herbivore (R^2^ = 0.98, 0.93), vertivore-herbivore (R^2^ = 0.99, 0.97), and invertivore-vertivore (R^2^ = 0.98, 0.90) analyses.

## Acknowledgments

We would like to thank Cole Bower, Maria Chikina, Allie Graham, Irene Kaplow, Emily Kopania, Johanna Kowalko, Greg Lang, Daniel Laverty, Linda Lowe-Krentz, Mary (Junjie) Ma, Kayleigh O’Keeffe, Andreas Pfenning, Matthew Pollard, Timothy Sackton, Abagael West, and members of the Meyer lab for consultation on the contents of the manuscript. We also thank two anonymous reviewers for their valuable feedback. We would like to thank Liam Revell for his assistance with the fossil-based ancestral reconstruction, Michael Hiller for providing the guide tree topology, and Derrick Um for writing the code to combine EltonTraits categories. We used OpenAI ChatGPT (5.6) and Google Gemini 3.6 for assistance writing simple code blocks. This material is based upon work supported by the National Science Foundation under grant no. 2233124 to W.K.M. Portions of this research were conducted on Lehigh University’s Research Computing infrastructure partially supported by NSF Award 2019035.

## Data Availability

The scripts used to complete this analysis, as well as the arguments used to execute them, can be found at https://github.com/wkmeyer-lab/RunRERPublication. A complete collection of the data used, including the input trees and alignment files, can be found at <u>Dryad</u>.

## References

Aizpurua O et al. 2021. The role of the gut microbiota in the dietary niche expansion of fishing bats. Anim Microbiome. 3:76. 10.1186/s42523-021-00137-w

Alhmoud JF, Woolley JF, Moustafa A-EA, Mallei MI. 2021. DNA Damage/Repair Management in Cancers. In: Advances in Medical Biochemistry, Genomics, Physiology, and Pathology. Jenny Stanford Publishing

Amador LI, Giannini NP. 2021. Evolution of diet in extant marsupials: emergent patterns from a broad phylogenetic perspective. Mammal Review. 51(2):178–192. 10.1111/mam.12223

Barts N et al. 2025. Gut bacterial and fungal communities of three rodent species respond uniquely to dietary fiber and protein manipulation. J Exp Biol. 228(14):jeb249794. 10.1242/jeb.249794

Baumgartner MR et al. 2014. Proposed guidelines for the diagnosis and management of methylmalonic and propionic acidemia. Orphanet J Rare Dis. 9(1):130. 10.1186/s13023-014-0130-8

Benjamini Y, Hochberg Y. 1995. Controlling the False Discovery Rate: A Practical and Powerful Approach to Multiple Testing. Royal Statistical Society Journal Series B: Methodological. 57(1):289–300. 10.1111/j.2517-6161.1995.tb02031.x

Bitar KN. 2003. Function of gastrointestinal smooth muscle: from signaling to contractile proteins. The American Journal of Medicine. 115(3, Supplement 1):15–23. 10.1016/S0002-9343(03)00189-X

Cambridge. 2025. [accessed 2025 Oct 13]. https://dictionary.cambridge.org/us/dictionary/english/carnivore

Cantalapiedra JL et al. 2014. Dietary innovations spurred the diversification of ruminants during the Caenozoic. Proceedings of the Royal Society B: Biological Sciences. 281(1776):20132746. 10.1098/rspb.2013.2746

Casotti G et al. 2006. Relationships between renal morphology and diet in 26 species of new world bats (suborder microchiroptera). Zoology. 109(3):196–207. 10.1016/j.zool.2006.03.003

Catania KC. 2013. Stereo and serial sniffing guide navigation to an odour source in a mammal. Nat Commun. 4(1):1441. 10.1038/ncomms2444

Catania KC. 2020. All in the Family – Touch Versus Olfaction in Moles. The Anatomical Record. 303(1):65–76. 10.1002/ar.24057

Chen M, Strömberg CAE, Wilson GP. 2019. Assembly of modern mammal community structure driven by Late Cretaceous dental evolution, rise of flowering plants, and dinosaur demise. Proceedings of the National Academy of Sciences. 116(20):9931–9940. 10.1073/pnas.1820863116

Chizzolini R, Zanardi E, Dorigoni V, Ghidini S. 1999. Calorific value and cholesterol content of normal and low-fat meat and meat products. Trends in Food Science & Technology. 10(4):119–128. 10.1016/S0924-2244(99)00034-5

Christmas MJ et al. 2023. Evolutionary constraint and innovation across hundreds of placental mammals. Science. 380(6643):eabn3943. 10.1126/science.abn3943

Chuy M, Alamargot D, Passerault J-M. 2012. Interference with prior knowledge while writing from sources: Effects of working memory capacity. Journal of Cognitive Psychology. 24(7):815–828. 10.1080/20445911.2012.699035

Coelho D et al. 2012. Mutations in ABCD4 cause a new inborn error of vitamin B12 metabolism. Nat Genet. 44(10):1152–1155. 10.1038/ng.2386

Compton ZT et al. 2025. Cancer Prevalence across Vertebrates. Cancer Discov. 15(1):227–244. 10.1158/2159-8290.CD-24-0573

Coogan SCP, Raubenheimer D, Stenhouse GB, Nielsen SE. 2014. Macronutrient optimization and seasonal diet mixing in a large omnivore, the grizzly bear: a geometric analysis. PLoS One. 9(5):e97968. 10.1371/journal.pone.0097968

Coombs EJ et al. 2022. The tempo of cetacean cranial evolution. Current Biology. 32(10):2233–2247.e4. 10.1016/j.cub.2022.04.060

Csárdi G et al. 2026. igraph: Network Analysis and Visualization. [accessed 2026 Oct 27]. https://cran.r-project.org/web/packages/igraph/index.html

Csardi G, Nepusz T. 2005. The Igraph Software Package for Complex Network Research. InterJournal. Complex Systems:1695

Das BK et al. 2025. Status of Ganges River dolphin Platanista gangetica (Lebeck, 1801) in the lower stretch of the Ganga River, India, with emphasis on threats, conservation, and recommendations. Front Ecol Evol. 13 [accessed 2026 Oct 27]. https://www.frontiersin.org/journals/ecology-and-evolution/articles/10.3389/fevo.2025.1523537/full. 10.3389/fevo.2025.1523537

Dierenfeld E, Alcorn H, Jacobsen K. 2002. [accessed 2025 Oct 13]. https://www.researchgate.net/publication/251886278_Nutrient_Composition_of_Whole_Vertebrate_Prey_Excluding_Fish_Fed_in_Zoos

Duque-Correa MJ et al. 2021. Mammalian intestinal allometry, phylogeny, trophic level and climate. Proc Biol Sci. 288(1944):20202888. 10.1098/rspb.2020.2888

Eisert R. 2011. Hypercarnivory and the brain: protein requirements of cats reconsidered. J Comp Physiol B. 181(1):1–17. 10.1007/s00360-010-0528-0

Emerling CA, Delsuc F, Nachman MW. 2018. Chitinase genes (CHIAs) provide genomic footprints of a post-Cretaceous dietary radiation in placental mammals. Science Advances. 4(5):eaar6478. 10.1126/sciadv.aar6478

Famoso NA, Hopkins SSB, Davis EB. 2018. How do diet and body mass drive reproductive strategies in mammals? Biol J Linn Soc. 124(2):151–156. 10.1093/biolinnean/bly038

Fruchterman TMJ, Reingold EM. 1991. Graph drawing by force-directed placement. Software: Practice and Experience. 21(11):1129–1164. 10.1002/spe.4380211102

Fyfe JC et al. 2004. The functional cobalamin (vitamin B12)–intrinsic factor receptor is a novel complex of cubilin and amnionless. Blood. 103(5):1573–1579. 10.1182/blood-2003-08-2852

The Gene Ontology Consortium. 2019. The Gene Ontology Resource: 20 years and still GOing strong. Nucleic Acids Res. 47(D1):D330–D338. 10.1093/nar/gky1055

Gill PG et al. 2014. Dietary specializations and diversity in feeding ecology of the earliest stem mammals. Nature. 512(7514):303–305. 10.1038/nature13622

Grant PR, Grant BR. 2006. Evolution of Character Displacement in Darwin’s Finches. Science. 313(5784):224–226. 10.1126/science.1128374

Grundler MC, Rabosky DL. 2021. Rapid increase in snake dietary diversity and complexity following the end-Cretaceous mass extinction. PLOS Biology. 19(10):e3001414. 10.1371/journal.pbio.3001414

GTEx. 2025a. [accessed 2025 Oct 28]. https://www.gtexportal.org/home/gene/CYP17A1

GTEx. 2025b. [accessed 2025 Oct 28]. https://www.gtexportal.org/home/gene/CYP21A2

GTEx. 2025c. [accessed 2025 Oct 28]. https://www.gtexportal.org/home/gene/HSD3B2

GTEx. 2025d. [accessed 2025 Oct 28]. https://www.gtexportal.org/home/gene/HSD11B2

GTEx Consortium. 2015. Human genomics. The Genotype-Tissue Expression (GTEx) pilot analysis: multitissue gene regulation in humans. Science. 348(6235):648–660. 10.1126/science.1262110

GTEx Portal. [accessed 2026a Aug 20]. https://www.gtexportal.org/home/gene/ENSG00000130948

GTEx Portal. [accessed 2026b Aug 20]. https://www.gtexportal.org/home/gene/ENSG00000086696

Hargreaves D, Buckland A, Sheaves M. 2017. Trophic guild concept: factors affecting within-guild consistency for tropical estuarine fish. Marine Ecology Progress Series. 564:175–186. 10.3354/meps11955

Hayden S et al. 2010. Ecological adaptation determines functional mammalian olfactory subgenomes. Genome Research. 20(1):1–9. 10.1101/gr.099416.109

He J et al. 2023. Piezo1 in Digestive System Function and Dysfunction. International Journal of Molecular Sciences. 24(16):12953. 10.3390/ijms241612953

Hecker N, Sharma V, Hiller M. 2019. Convergent gene losses illuminate metabolic and physiological changes in herbivores and carnivores. Proceedings of the National Academy of Sciences. 116(8):3036–3041. 10.1073/pnas.1818504116

Hiller M. 2024. Personal Communication.

Hillier SG. 2017. On gonads and gadflies: the estrus angle. J Endocrinol. 233(3):C1–C8. 10.1530/JOE-17-0136

Hu Y et al. 2017. Comparative genomics reveals convergent evolution between the bamboo-eating giant and red pandas. Proceedings of the National Academy of Sciences. 114(5):1081–1086. 10.1073/pnas.1613870114

Hughes GM et al. 2018. The Birth and Death of Olfactory Receptor Gene Families in Mammalian Niche Adaptation. Mol Biol Evol. 35(6):1390–1406. 10.1093/molbev/msy028

Iason GR, Villalba JJ. 2006. Behavioral Strategies of Mammal Herbivores Against Plant Secondary Metabolites: The Avoidance–Tolerance Continuum. J Chem Ecol. 32(6):1115–1132. 10.1007/s10886-006-9075-2

Izadi S, Aguilar de Soto N, Constantine R, Johnson M. 2022. Feeding tactics of resident Bryde’s whales in New Zealand. Marine Mammal Science. 38(3):1104–1117. 10.1111/mms.12918

Jain A, Tuteja G. 2019. TissueEnrich: Tissue-specific gene enrichment analysis. Bioinformatics. 35(11):1966–1967. 10.1093/bioinformatics/bty890

Jones CA, Reynolds SE. 1980. A reinvestigation of the effects of cortisol on growth in insects. Journal of Insect Physiology. 26(9):601–605. 10.1016/0022-1910(80)90028-1

Jones KE et al. 2009. PanTHERIA: a species-level database of life history, ecology, and geography of extant and recently extinct mammals. Ecology. 90(9):2648–2648. 10.1890/08-1494.1

Kambere MB, Lane RP. 2007. Co-regulation of a large and rapidly evolving repertoire of odorant receptor genes. BMC Neurosci. 8(3):S2. 10.1186/1471-2202-8-S3-S2

Kanehisa M, Goto S. 2000. KEGG: kyoto encyclopedia of genes and genomes. Nucleic Acids Res. 28(1):27–30. 10.1093/nar/28.1.27

Karasov WH, Rio CM del, Caviedes-Vidal E. 2011. Ecological Physiology of Diet and Digestive Systems. Annual Review of Physiology. 73(Volume 73, 2011):69–93. 10.1146/annurev-physiol-012110-142152

Kelley NP, Motani R. 2015. Trophic convergence drives morphological convergence in marine tetrapods. Biology Letters. 11(1):20140709. 10.1098/rsbl.2014.0709

Kim Soonok et al. 2016. Comparison of carnivore, omnivore, and herbivore mammalian genomes with a new leopard assembly. Genome Biology. 17(1):211. 10.1186/s13059-016-1071-4

Kirilenko BM, et al. 2023. Integrating gene annotation with orthology inference at scale. Science. 380(6643):eabn3107 [accessed 2024 July 23]. https://www.science.org/doi/10.1126/science.abn3107. 10.1126/science.abn3107

Kissling WD et al. 2014. Establishing macroecological trait datasets: digitalization, extrapolation, and validation of diet preferences in terrestrial mammals worldwide. Ecology and Evolution. 4(14):2913–2930. 10.1002/ece3.1136

Kowalczyk A et al. 2019. RERconverge: An R package for associating evolutionary rates with convergent traits. Bioinformatics. 35(22):4815–4817 [accessed 2020 Aug 25]. https://academic.oup.com/bioinformatics/article/35/22/4815/5514536. 10.1093/bioinformatics/btz468

Kowalczyk A, Partha R, Clark NL, Chikina M. 2020. Pan-mammalian analysis of molecular constraints underlying extended lifespan. eLife. 9:e51089. 10.7554/eLife.51089

Le SQ, Gascuel O. 2008. An Improved General Amino Acid Replacement Matrix. Mol Biol Evol. 25(7):1307–1320. 10.1093/molbev/msn067

Li D, Zhang J. 2014. Diet Shapes the Evolution of the Vertebrate Bitter Taste Receptor Gene Repertoire. Mol Biol Evol. 31(2):303–309. 10.1093/molbev/mst219

Liberzon A et al. 2011. Molecular signatures database (MSigDB) 3.0. Bioinformatics. 27(12):1739–1740. 10.1093/bioinformatics/btr260

Lintulaakso K, Tatti N, Žliobaitė I. 2023. Quantifying mammalian diets. Mamm Biol. 103(1):53–67. 10.1007/s42991-022-00323-6

Louca S, Doebeli M. 2018. Efficient comparative phylogenetics on large trees. Bioinformatics. 34(6):1053–1055 [accessed 2025 July 13]. 10.1093/bioinformatics/btx701. 10.1093/bioinformatics/btx701

Lustig C, May CP, Hasher L. 2001. Working memory span and the role of proactive interference. Journal of Experimental Psychology: General. 130(2):199–207. 10.1037/0096-3445.130.2.199

Ma J et al. 2023. Gene expressions between obligate bamboo-eating pandas and non-herbivorous mammals reveal converged specialized bamboo diet adaptation. BMC Genomics. 24(1):23. 10.1186/s12864-023-09111-z

Macdonald AR, James ME, Mitchell JD, Holland BR. 2025. From Trees to Traits: A Review of Advances in PhyloG2P Methods and Future Directions. Genome Biol Evol. 17(9):evaf150. 10.1093/gbe/evaf150

Machovsky-Capuska GE, Senior AM, Simpson SJ, Raubenheimer D. 2016. The Multidimensional Nutritional Niche. Trends in Ecology & Evolution. 31(5):355–365. 10.1016/j.tree.2016.02.009

de Magalhães JP et al. 2024. Human Ageing Genomic Resources: updates on key databases in ageing research. Nucleic Acids Res. 52(D1):D900–D908. 10.1093/nar/gkad927

Malmberg JL, White LA, VandeWoude S. 2021. Bioaccumulation of Pathogen Exposure in Top Predators. Trends in Ecology & Evolution. 36(5):411–420. 10.1016/j.tree.2021.01.008

de Matos Dias D, de Campos CB, Guimarães Rodrigues FH. 2018. Behavioural ecology in a predator-prey system. Mamm Biol. 92(1):30–36. 10.1016/j.mambio.2018.04.005

Mayani-Parás F et al. 2023. Classification and distribution of functional groups of birds and mammals in Mexico. PLoS One. 18(11):e0287036. 10.1371/journal.pone.0287036

Meloro C, Hudson A, Rook L. 2015. Feeding habits of extant and fossil canids as determined by their skull geometry. Journal of Zoology. 295(3):178–188. 10.1111/jzo.12196

Merriam-Webster. 2025. [accessed 2025 Oct 13]. https://www.merriam-webster.com/dictionary/carnivore

Milacic M et al. 2024. The Reactome Pathway Knowledgebase 2024. Nucleic Acids Res. 52(D1):D672–D678. 10.1093/nar/gkad1025

Miller CV, Pittman M. 2021. The diet of early birds based on modern and fossil evidence and a new framework for its reconstruction. Biological Reviews. 96(5):2058–2112. 10.1111/brv.12743

Monteiro LR, Nogueira MR. 2011. Evolutionary patterns and processes in the radiation of phyllostomid bats. BMC Evol Biol. 11(1):137. 10.1186/1471-2148-11-137

Mu Y et al. 2021. Molecular Evolution of Tooth-Related Genes Provides New Insights into Dietary Adaptations of Mammals. J Mol Evol. 89(7):458–471. 10.1007/s00239-021-10017-1

Munoz-Lopez M, Garcia-Perez JL. 2012. DNA Transposons: Nature and Applications in Genomics. Current Genomics. 11(2):115–128. 10.2174/138920210790886871

Nijhout HF, Laub E. 2018. The role of hormones. In: Córdoba-Aguilar A, González-Tokman D, González-Santoyo I, editors. Insect Behavior: From Mechanisms to Ecological and Evolutionary Consequences. Oxford University Press; p 0 [accessed 2025 Dec 4]. 10.1093/oso/9780198797500.003.0004. 10.1093/oso/9780198797500.003.0004

Novick P, Smith J, Ray D, Boissinot S. 2010. Independent and parallel lateral transfer of DNA transposons in tetrapod genomes. Gene. 449(1):85–94. 10.1016/j.gene.2009.08.017

Nowak RM. 1999. Walker’s Mammals of the World. Johns Hopkins University Press. [accessed 2026 June 19]. https://www.press.jhu.edu/books/title/2993/walkers-mammals-world. 10.56021/9780801857898

Ocampo M, Pincheira-Donoso D, Sayol F, Rios RS. 2022. Evolutionary transitions in diet influence the exceptional diversification of a lizard adaptive radiation. BMC Ecol Evo. 22(1):74. 10.1186/s12862-022-02028-3

Osmanski AB et al. 2023. Insights into mammalian TE diversity through the curation of 248 genome assemblies. Science. 380(6643):eabn1430. 10.1126/science.abn1430

Oxford Reference. Oxford Reference; [accessed 2025 Oct 13]. https://www.oxfordreference.com/display/10.1093/oi/authority.20110803095550817. 10.1093/oi/authority.20110803095550817

Pandelis GG, Grundler MC, Rabosky DL. 2023. Ecological correlates of cranial evolution in the megaradiation of dipsadine snakes. BMC Ecol Evol. 23:48. 10.1186/s12862-023-02157-3

Paradis E, Schliep K. 2019. ape 5.0: an environment for modern phylogenetics and evolutionary analyses in R. Bioinformatics. 35(3):526–528 [accessed 2026 June 19]. 10.1093/bioinformatics/bty633. 10.1093/bioinformatics/bty633

Partha R, Kowalczyk A, Clark NL, Chikina M. 2019. Robust Method for Detecting Convergent Shifts in Evolutionary Rates. Mol Biol Evol. 36(8):1817–1830. 10.1093/molbev/msz107

Pedersen TL, RStudio. 2025. ggraph: An Implementation of Grammar of Graphics for Graphs and Networks. [accessed 2026 Oct 27]. https://cran.r-project.org/web/packages/ggraph/index.html

Phan A, Joshi P, Kadelka C, Friedberg I. 2025. A longitudinal analysis of function annotations of the human proteome reveals consistently high biases. Database (Oxford). 2025:baaf036. 10.1093/database/baaf036

Pineda-Munoz S, Alroy J. 2014. Dietary characterization of terrestrial mammals. Proceedings of the Royal Society B: Biological Sciences. 281(1789):20141173. 10.1098/rspb.2014.1173

Piñero J et al. 2017. DisGeNET: a comprehensive platform integrating information on human disease-associated genes and variants. Nucleic Acids Res. 45(D1):D833–D839. 10.1093/nar/gkw943

Pollard MD, Meyer WK, Puckett EE. 2024. Convergent relaxation of molecular constraint in herbivores reveals the changing role of liver and kidney functions across mammalian diets. Genome Res. 34(12):2176–2189. 10.1101/gr.278930.124

Pond SLK, Frost SDW, Muse SV. 2005. HyPhy: hypothesis testing using phylogenies. Bioinformatics. 21(5):676–679. 10.1093/bioinformatics/bti079

Price SA, Hopkins SSB. 2015. The macroevolutionary relationship between diet and body mass across mammals. Biological Journal of the Linnean Society. 115(1):173–184. 10.1111/bij.12495

Price SA, Hopkins SSB, Smith KK, Roth VL. 2012. Tempo of trophic evolution and its impact on mammalian diversification. Proceedings of the National Academy of Sciences. 109(18):7008–7012. 10.1073/pnas.1117133109

PubChem. 2025. [accessed 2025 Oct 28]. https://pubchem.ncbi.nlm.nih.gov/gene/HSD11B1/human

PubChem. [accessed 2026a Oct 27]. https://pubchem.ncbi.nlm.nih.gov/gene/HSD17B2/human

PubChem. [accessed 2026b Oct 27]. https://pubchem.ncbi.nlm.nih.gov/gene/HSD17B3/human

Redlich R et al. 2024. RERconverge Expansion: Using Relative Evolutionary Rates to Study Complex Categorical Trait Evolution. Molecular Biology and Evolution. 41(11):msae210 [accessed 2024 Dec 23]. 10.1093/molbev/msae210. 10.1093/molbev/msae210

Reuter DM, Hopkins SSB, Price SA. 2023. What is a mammalian omnivore? Insights into terrestrial mammalian diet diversity, body mass and evolution. Proceedings of the Royal Society B: Biological Sciences. 290(1992):20221062. 10.1098/rspb.2022.1062

Revell L. 2025. [accessed 2025 Dec 10]. https://blog.phytools.org/2025/09/ancestral-states-for-discrete-trait.html

Revell LJ. 2012. phytools: an R package for phylogenetic comparative biology (and other things). Methods in Ecology and Evolution. 3(2):217–223 [accessed 2026 June 19]. https://onlinelibrary.wiley.com/doi/abs/10.1111/j.2041-210X.2011.00169.x. 10.1111/j.2041-210X.2011.00169.x

Román-Palacios C, Scholl JP, Wiens JJ. 2019. Evolution of diet across the animal tree of life. Evol Lett. 3(4):339–347. 10.1002/evl3.127

Rothman JM et al. 2014. Nutritional contributions of insects to primate diets: Implications for primate evolution. Journal of Human Evolution. 71:59–69 (The Other Faunivory: The Significance of Insects & Insect Resources for Nonhuman Primates, Modern Humans, & Extinct Hominins). 10.1016/j.jhevol.2014.02.016

Sadier A et al. 2021. Making a bat: The developmental basis of bat evolution. Genet Mol Biol. 43(1 Suppl 2):e20190146. 10.1590/1678-4685-GMB-2019-0146

Sapolsky RM, Romero LM, Munck AU. 2000. How do glucocorticoids influence stress responses? Integrating permissive, suppressive, stimulatory, and preparative actions. Endocr Rev. 21(1):55–89. 10.1210/edrv.21.1.0389

Saputra E et al. 2021. Phylogenetic Permulations: A Statistically Rigorous Approach to Measure Confidence in Associations in a Phylogenetic Context. Mol Biol Evol. 38(7):3004–3021. 10.1093/molbev/msab068

Schliep KP. 2011. phangorn: Phylogenetic analysis in R. Bioinformatics. 27(4):592–593 [accessed 2019 Sept 18]. https://academic.oup.com/bioinformatics/article-lookup/doi/10.1093/bioinformatics/btq706. 10.1093/bioinformatics/btq706

Senti T, Gifford M. 2024. Seasonal and taxonomic variation in arthropod macronutrient content. Food Webs. 38:e00328. 10.1016/j.fooweb.2023.e00328

Shtulman A, Valcarcel J. 2012. Scientific knowledge suppresses but does not supplant earlier intuitions. Cognition. 124(2):209–215. 10.1016/j.cognition.2012.04.005

Slater GJ, Friscia AR. 2019. Hierarchy in adaptive radiation: A case study using the Carnivora (Mammalia). Evol. 73(3):524–539. 10.1111/evo.13689

Smith CL, Goldsmith C-AW, Eppig JT. 2004. The Mammalian Phenotype Ontology as a tool for annotating, analyzing and comparing phenotypic information. Genome Biology. 6(1):R7. 10.1186/gb-2004-6-1-r7

Smith SD, Pennell MW, Dunn CW, Edwards SV. 2020. Phylogenetics is the New Genetics (for Most of Biodiversity). Trends in Ecology & Evolution. 35(5):415–425. 10.1016/j.tree.2020.01.005

Stelzer G et al. 2016. The GeneCards Suite: From Gene Data Mining to Disease Genome Sequence Analyses. Curr Protoc Bioinformatics. 54:1.30.1-1.30.33. 10.1002/cpbi.5

Stephens RB, Ouimette AP, Hobbie EA, Rowe RJ. 2022. Reevaluating trophic discrimination factors (Δδ13C and Δδ15N) for diet reconstruction. Ecological Monographs. 92(3):e1525. 10.1002/ecm.1525

Stricker J et al. 2021. Interference between naïve and scientific theories occurs in mathematics and is related to mathematical achievement. Cognition. 214:104789. 10.1016/j.cognition.2021.104789

Subramanian A et al. 2005. Gene set enrichment analysis: A knowledge-based approach for interpreting genome-wide expression profiles. Proceedings of the National Academy of Sciences. 102(43):15545–15550. 10.1073/pnas.0506580102

Wagner F et al. 2022. Reconstruction of evolutionary changes in fat and toxin consumption reveals associations with gene losses in mammals: A case study for the lipase inhibitor PNLIPRP1 and the xenobiotic receptor NR1I3. Journal of Evolutionary Biology. 35(2):225–239. 10.1111/jeb.13970

Wang Z et al. 2016. Evolution of Digestive Enzymes and RNASE1 Provides Insights into Dietary Switch of Cetaceans. Mol Biol Evol. 33(12):3144–3157. 10.1093/molbev/msw191

Watkins D, Rosenblatt DS. 2011. Inborn errors of cobalamin absorption and metabolism. American Journal of Medical Genetics Part C: Seminars in Medical Genetics. 157(1):33–44. 10.1002/ajmg.c.30288

Wertheim JO et al. 2015. RELAX: Detecting Relaxed Selection in a Phylogenetic Framework. Mol Biol Evol. 32(3):820–832. 10.1093/molbev/msu400

Wilhoit K et al. 2025. Convergent Evolution and Predictability of Gene Copy Numbers Associated with Diets in Mammals. Genome Biol Evol. 17(2):evaf008. 10.1093/gbe/evaf008

Wilman H et al. 2014. EltonTraits 1.0: Species-level foraging attributes of the world’s birds and mammals. Ecology. 95(7):2027–2027 [accessed 2022 June 1]. https://onlinelibrary.wiley.com/doi/full/10.1890/13-1917.1. 10.1890/13-1917.1

Wirthlin M et al. 2019. A Modular Approach to Vocal Learning: Disentangling the Diversity of a Complex Behavioral Trait. Neuron. 104(1):87–99. 10.1016/j.neuron.2019.09.036

Wirthlin ME et al. 2024. Vocal learning-associated convergent evolution in mammalian proteins and regulatory elements. Science. 0(0):eabn3263 [accessed 2024 Mar 22]. https://www.science.org/doi/10.1126/science.abn3263. 10.1126/science.abn3263

Wu Y. 2022. Diet evolution of carnivorous and herbivorous mammals in Laurasiatheria. BMC Ecol Evo. 22(1):82. 10.1186/s12862-022-02033-6

Zhukova NV. 2019. Fatty Acids of Marine Mollusks: Impact of Diet, Bacterial Symbiosis and Biosynthetic Potential. Biomolecules. 9(12):857. 10.3390/biom9120857

