## Supplementary Text for "Diet-Related Molecular Evolution Differs among Vertivores, Invertivores, and Combined Carnivores"

#### Comparison of overlapping K-R categories

We see a similar distribution of overlap in K-R categories between analyses as we see with individual genes (Figure 3). 88 (80%) of the 110 K-R categories significant in the combined carnivore-herbivore analysis were also significant in the invertivore-herbivore analysis, but only 3 (3%) were significant in the vertivore-herbivore analysis (Supplementary Figure 1). We see this pattern in both the gene sets evolving faster in herbivores than predatory diets (Figure S1B), and in gene sets evolving faster in predatory diets than herbivores (Figure S1C).

However, direct comparisons of numbers of significant gene set categories are known to have issues. One consideration is that the annotations of gene sets are biased toward particular functions and processes (Phan et al. 2025). Additionally, gene sets often have high (and highly variable) levels of overlap. These biases can mean that an increased number of significant gene set categories can be a result of enrichment of processes that are more finely annotated, rather than an increase in the distinct biological processes that are enriched. Relevant to our case is the high level of annotation of signaling pathways (Phan et al. 2025), as signaling is one of the processes that is enriched for genes evolving faster in species with predatory diets than in herbivores. This is one of the drivers of a greater number of significant gene sets evolving faster in predators than in herbivores, despite fewer individually significant genes that are evolving faster in predators than in herbivores.

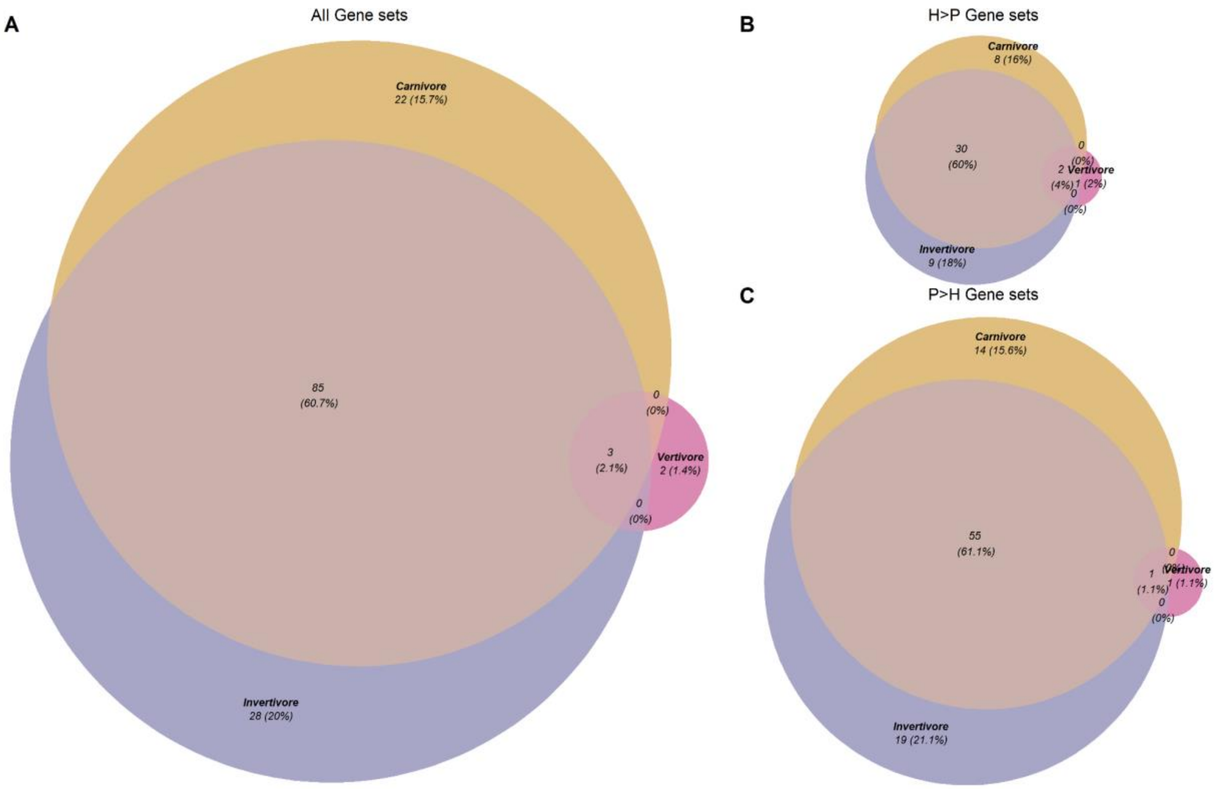


Supplementary Figure 1: Representations of the overlap across analyses comparing K-R categories enriched for genes with differences in relative evolutionary rates (RER) between herbivores and lineages with predatory diets. Categories included have FDR-adjusted P <0.05 and are robust to species set selection. Venn diagrams of (A) all categories with robust significantly different RER in each predatory diet relative to herbivory, (B) categories with higher RER in herbivores than in species with predatory diets, and (C) categories with higher RER in species with predatory diets than in herbivores. Circle sizes represent the same scale throughout the figure. Note that percentages in the figure represent percentages of categories in each Venn diagram, while percentages in the text represent percentages of genes for individual diets.

### Exploration of phenotypes potentially underlying enrichment of DNA repair pathways

We hypothesized that the findings of DNA repair genes evolving more slowly in invertivores than in vertivores or herbivores might be due to another phenotype that covaries with diet in our dataset. To assess this possibility, we tested for differences in the distribution of three continuous phenotypes with the potential to influence selection on DNA repair genes –

maximum lifespan, neoplasia (tumor) prevalence, and young DNA transposon proportion – across diets. Lifespan has been previously linked with DNA repair, and failures of DNA repair is a common cause of cancer (Munoz-Lopez and Garcia-Perez 2012; Kowalczyk et al. 2020; Alhmoud et al. 2021). We first identified overlapping species between our dataset and those of three prior studies (Osmanski et al. 2023; de Magalhães et al. 2024; Compton et al. 2025) (Supplementary Table 1). For each phenotype, we then used the edgeVars function included within the RERconverge package to infer ancestral mean values and changes  along branches within a tree including the overlapping species. We then tested for a difference in the distribution of either mean phenotype or phenotype change across branches with different diets from our four diet analyses, using a Kruskal-Wallis test followed by a post hoc Dunn's Z test for each phenotype.

While we found differences in distributions of the inferred means of all three phenotypes – maximum lifespan (Kruskal-Wallis $\chi^2$ = 13.3, P = 4.0 x $10^{-3}$), neoplasia (tumor) prevalence ($\chi^2$ = 12.9, P = 4.9 x $10^{-3}$), and young DNA transposon proportion ($\chi^2$ = 32.7, P = 3.8 x $10^{-7}$) (Supplementary Figure 2), only in the case of young DNA transposon proportion did the post hoc tests indicate that these differences were driven predominantly by the distribution for invertivores (Dunn's Z = 4.1 for the invertivore-vertivore comparison, P = 4.0 x $10^{-5}$) (Supplementary Table 9). This is the pattern predicted for a phenotype that contributes to the observed enrichment of DNA repair genes among genes evolving slowly in invertivores in our dataset. We thus performed additional analyses to assess whether the relative evolutionary rates of 9 DNA repair genes (chosen due to high ranks within enriched categories in our invertivore-vertivore comparisons) were correlated with inferred means or changes in transposon content across branches of the tree, using the continuous version of RERconverge with default settings (Kowalczyk et al. 2019; Partha et al. 2019). We found significant correlations between mean transposon content and relative evolutionary rates for several DNA repair genes (e.g., *EME2*, *USP45*), and all 9 genes that we tested had negative correlations (Supplementary Table 10). Taken together, this suggests that variation in DNA transposon content across diets could contribute to the signature of enrichment of DNA repair genes among genes evolving slowly in invertivores.

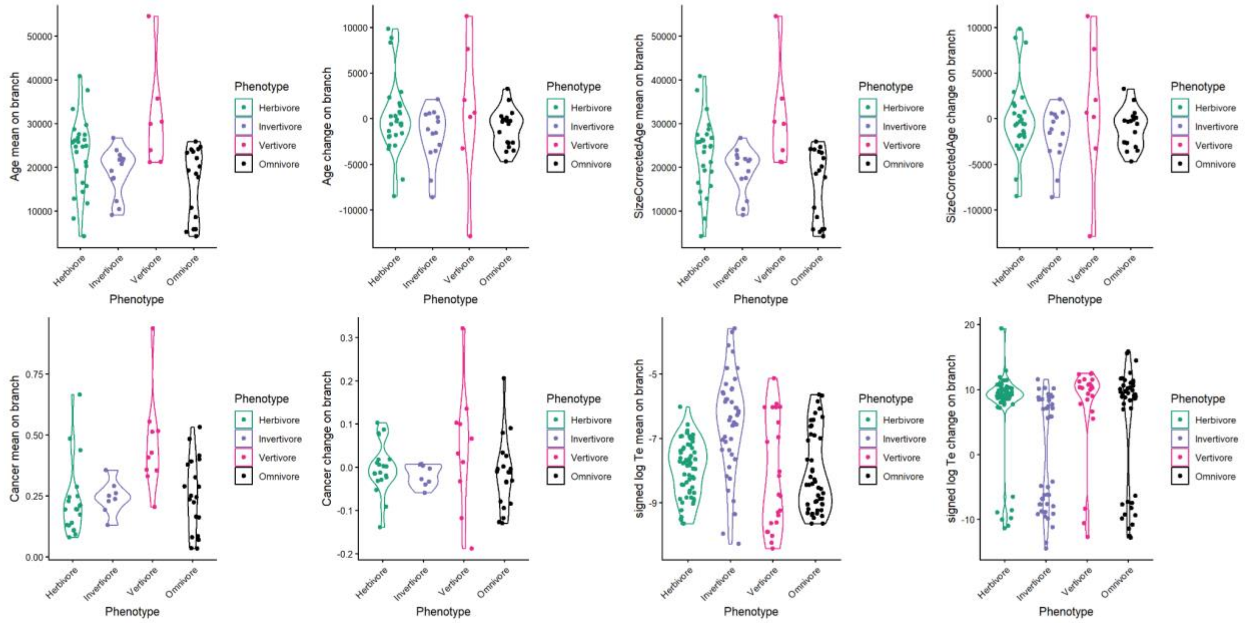


Supplementary Figure 2: Distributions of inferred means and changes in four continuous phenotypes that could be relevant to the evolution of DNA repair genes, across lineages with four diets. Mean on branch = mean value of the phenotype on each branch inferred using RERconverge ‘char2Paths’ with ‘metric = “mean”’. Change on branch = change in the phenotype on each branch inferred using RERconverge ‘char2Paths’ with ‘metric = “diff”’. Age = maximum lifespan in days (Jones et al. 2009; de Magalhães et al. 2024)(ref). Size-corrected age = Residuals of a regression of maximum lifespan (in days) on body mass (in grams) (Jones et al. 2009; de Magalhães et al. 2024). Cancer = neoplasia prevalence (Compton et al. 2025). Signed log Te = signed log(young DNA proportion)(Osmanski et al. 2023).

#### K-R categories evolving slower in herbivores than predatory diets

As described in the results, we found a large number of K-R categories which are evolving slower in herbivores than the various predatory diets. We chose to include the graph of these genes sets in the supplement for length. Category names for these pathways can be found in Supplementary Table 7.

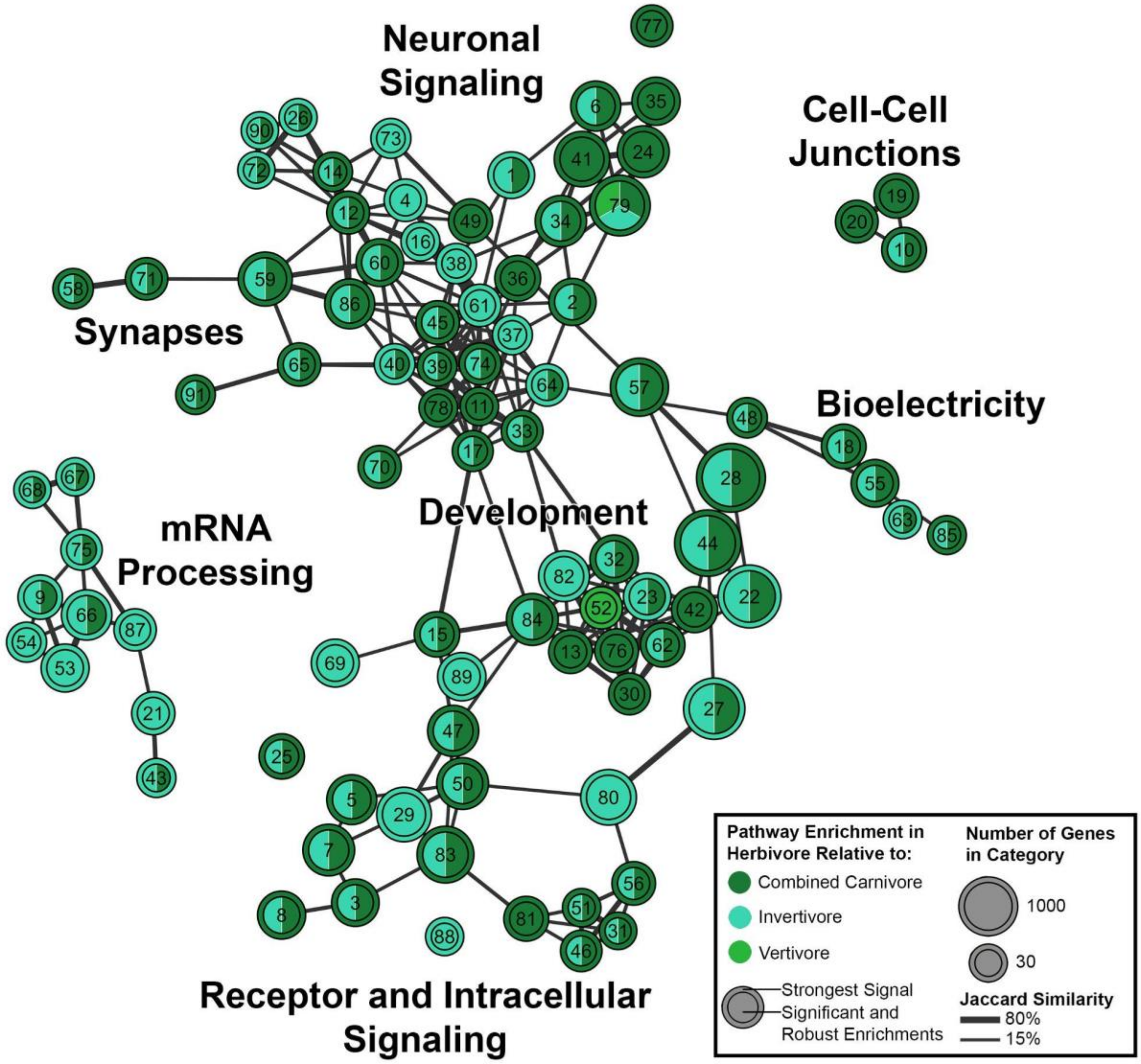


Supplementary Figure 3: Network graph of K-R categories enriched for genes evolving more slowly in species with herbivorous diets than in predatory species, with at least one such enrichment significant at FDR = 0.05 and robust to species selection. Clusters were generated using the igraph R package (Csardi and Nepusz 2005) and labeled manually based on general function. Each node represents a K-R category; see Supplementary Table 7 for individual category names. Nodes are connected by lines with thickness representing the Jaccard similarity. Color within the circle indicates for which comparison(s) we observe significant enrichment (dark green: combined carnivore-herbivore; teal: invertivore-herbivore; light green: vertivore-herbivore). Circle outlines indicate the comparison with the strongest signal from the RER analysis.

#### Analysis of alternate species sets

We found that most of the genes found to be significant (FDR-adjusted $P < 0.05$) in our main analyses were significant at this level in a large number of the alternates as well. For the invertivore-vertivore comparison, genes significant in the main analysis were significant in an average of 76 of the 100 alternate species sets, and 86% of the significant genes from the main analysis were significant in ≥50% of the alternate species sets (Supplementary Figure 3A; Supplementary Table 2). The mean number of significant alternates for a significant main analysis gene is lower in the other diet comparisons, though still fairly high (combined carnivore-herbivore 56/100, invertivore-herbivore 64/100, vertivore-herbivore 48/100). The percentage of main analysis significant genes robust to alternates is also lower (combined carnivore-herbivore 43%, invertivore-herbivore 33%, vertivore-herbivore 54%) (Supplementary Figure 3A).

A similar pattern can be seen in the K-R category robustness, with the invertivore-vertivore analysis having the by far the highest robustness (mean alternates of significant 86/100, 87% of results robust), and the other comparisons being less robust, with the vertivore-herbivore comparisons being particularly affected (combined carnivore-herbivore 42/100, 39%; invertivore-herbivore 53/100, 54%; vertivore-herbivore 20/100 15%) (Supplementary Figure 3B).

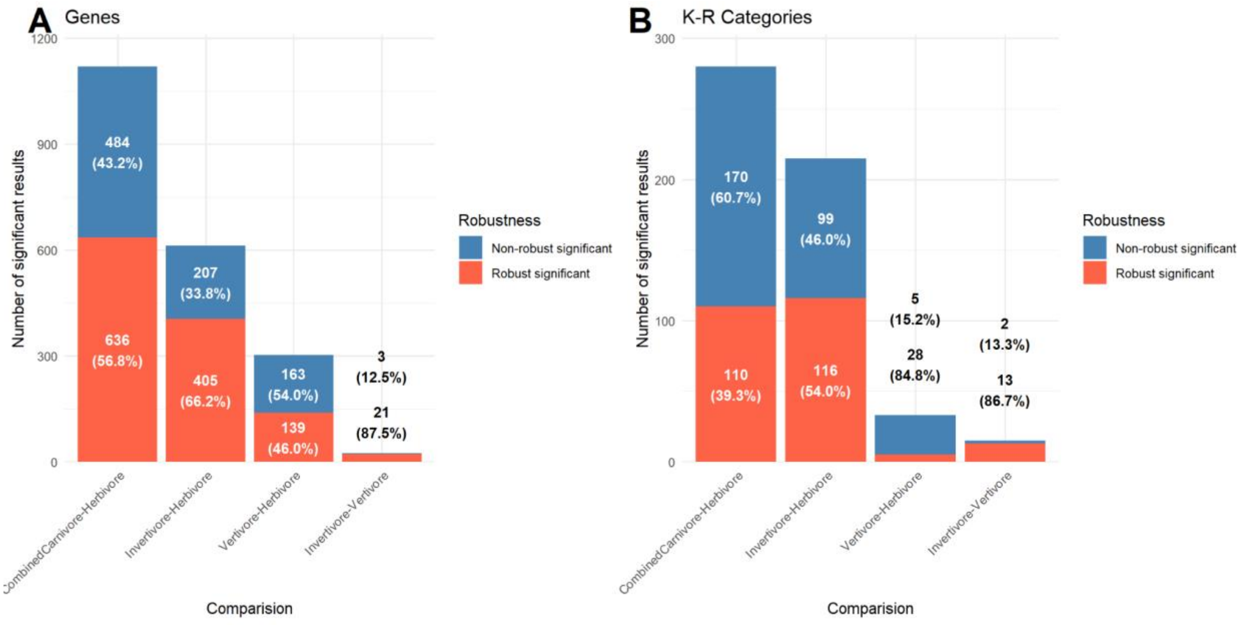


Supplementary Figure 4: Bar plots of the ratio of results significant in the main analysis which are robust to (significant in ≥50%) alternative species selection. Bar heights are the total number of significant results, with the orange section being robust and the blue section being not robust, with (A) displaying significant genes and (B) displaying significant K-R categories.

#### Strict Four-Category Diet Definition

In addition to the main analysis, we performed a separate analysis using a 90% threshold for all of the four diet categories, to assess the robustness of our results to different decisions in the assignment of combined carnivores to the two nuanced predatory diets. In this strict analysis, species with diets consisting of ≥90% herbivorous categories were considered herbivores; ≥90% vertivorous categories vertivores; ≥90% invertivorous categories invertivores. Species with diets consisting of ≥90% (vertivorous categories + invertivorous categories), but not ≥90%

vertivorous categories or ≥90% invertivorous categories were considered ‘mixed predators’, neither vertivores or invertivores. These mixed predators were dropped from the four diet analysis, resulting in a different species set between the three diet and four diet analyses.

The strength of signal for genes are highly correlated between this stricter definition and the main definition in the combined carnivore-herbivore ($R^2$=0.88), invertivore-herbivore ($R^2$=0.91), vertivore-herbivore ($R^2$=0.72), and invertivore-vertivore ($R^2$=0.73) analyses (Supplementary Figure 4). It is also very strong for K-R categories in the combined carnivore-herbivore ($R^2$=0.81) and invertivore-herbivore ($R^2$=0.85) analyses, and fairly strong in the vertivore-herbivore ($R^2$=0.44) and invertivore-vertivore ($R^2$= 0.59) analyses.

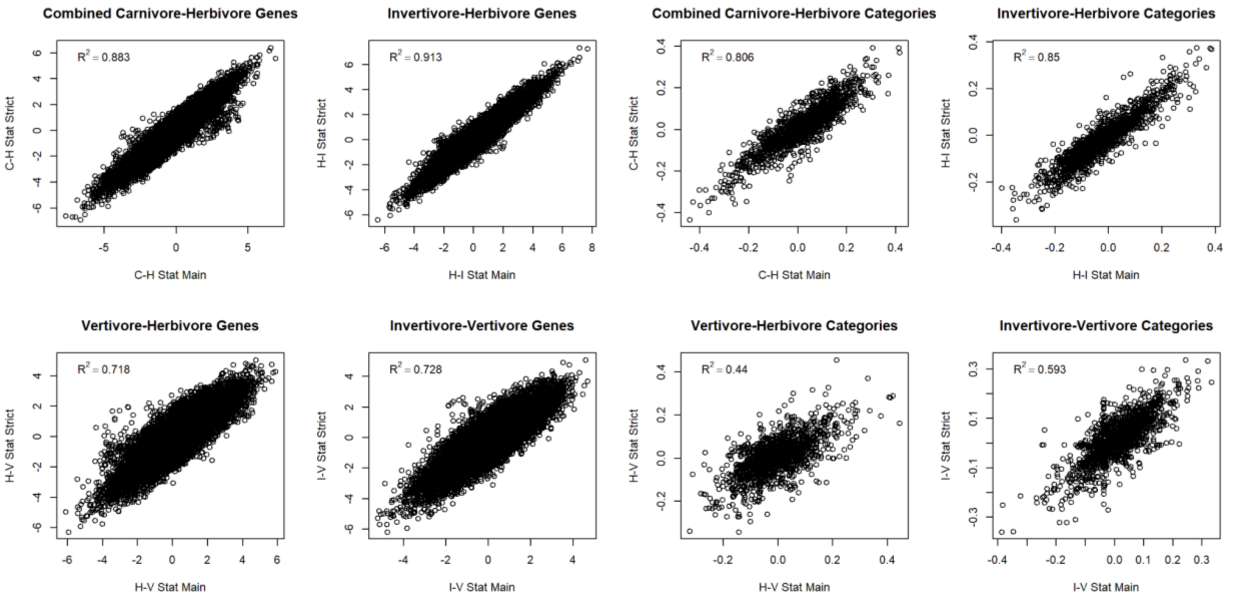


Supplementary Figure 5: Correlation of stat (genes: Dunn Z; K-R categories Mann Whitney U AUC) of main analysis results with results in the strict nuanced predator analysis .

### Permulations Analyses

We found the results of our main analyses to be mostly in concordance with the results from the permulation analyses. The log permulation-based P-values, signed by direction of effect, across all genes were strongly correlated with the main signed log P-values for each pairwise post hoc comparison involving at least one predatory diet ($r_s$: 0.88 herbivore-invertivore, 0.84 herbivore-vertivore, 0.81 invertivore-vertivore) (Supplementary Figure 5). We found that all of the robust significantly enriched (FDR < 0.05 in ≥50% of alternates) K-R categories in the invertivore-vertivore, invertivore-herbivore, and vertivore-herbivore post hoc analyses were also significantly enriched in the corresponding permulation analyses.

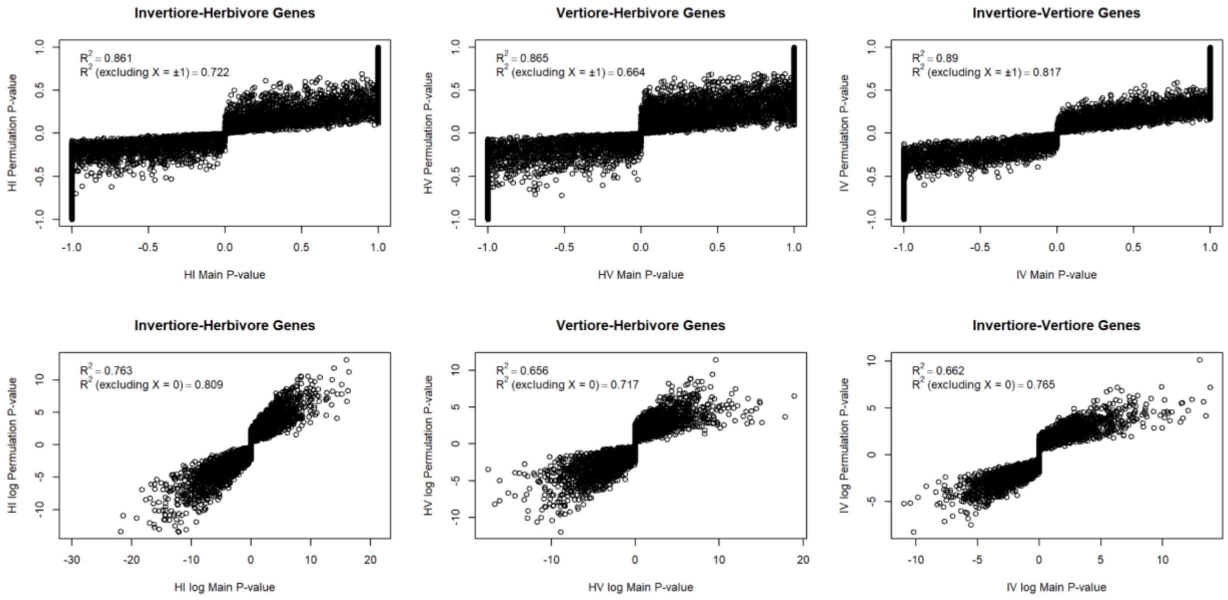


Supplementary Figure 6: Correlation of gene direction-of-effect (sign of Dunn Z score) signed P-values and log P-values between the main analysis and permulation analysis. $R^2$ values included for both the entire dataset, and correlations which exclude genes with a P-value of 1 in the main analysis.

#### Additional gene set enrichment analyses

We performed gene set enrichment analysis using the MGI_Mammalian_Phenotype_Level_4, GO_Biological_Process_2023, and MSigDb's C2 gene sets (Smith et al. 2004; Liberzon et al. 2011; Piñero et al. 2017; The Gene Ontology Consortium 2019). While these results are not discussed in the manuscript, they can be found in Supplementary Data 3. Similarly, we performed but do not discuss enrichment analysis on gene sets based on tissue of expression. These are based on the GTEx Consortium data, with a gene being considered tissue specific if it was expressed at least five-fold higher in up to seven tissues compared to all others (GTEx Consortium 2015; Jain and Tuteja 2019; Pollard et al. 2024). These results can be found in Supplementary Data 4.
